# Molecular encoding of stimulus features in a single sensory neuron type enables neuronal and behavioral plasticity

**DOI:** 10.1101/2023.01.22.525070

**Authors:** Nathan Harris, Samuel Bates, Zihao Zhuang, Matthew Bernstein, Jamie Stonemetz, Tyler Hill, Yanxun V. Yu, John A. Calarco, Piali Sengupta

## Abstract

Neurons modify their transcriptomes in response to an animal’s experience. How specific experiences are transduced to modulate gene expression and precisely tune neuronal functions are not fully defined. Here, we describe the molecular profile of a thermosensory neuron pair in *C. elegans* experiencing different temperature stimuli. We find that distinct salient features of the temperature stimulus including its duration, magnitude of change, and absolute value are encoded in the gene expression program in this single neuron, and identify a novel transmembrane protein and a transcription factor whose specific transcriptional dynamics are essential to drive neuronal, behavioral, and developmental plasticity. Expression changes are driven by broadly expressed activity-dependent transcription factors and corresponding *cis*-regulatory elements that nevertheless direct neuron- and stimulus-specific gene expression programs. Our results indicate that coupling of defined stimulus characteristics to the gene regulatory logic in individual specialized neuron types can customize neuronal properties to drive precise behavioral adaptation.

## Introduction

Animals must modulate their physiology and behavior appropriately in response to changing environmental conditions. A major driving force underlying long-term plasticity is the ability of cells to reformat the usage of their genomes as a function of their experience (Lee et al., 2008; Roselli et al., 2021; Bliim et al., 2016; Yap and Greenberg 2018). In the nervous system, different patterns of activity are translated into unique gene expression programs across neuronal ensembles comprised of multiple neuronal subtypes (Spiegel et al., 2014; Hu et al., 2017; Tyssowski et al., 2018; Hrvatin et al., 2018; Lacar et al., 2016; Wu et al., 2017). However, since neurons exhibit extensive heterogeneity in their properties even within a defined subtype (Peng et al., 2021; Zeng, 2022; Scala et al., 2021; Cembrowski and Spruston, 2019), whether behavioral plasticity in response to a defined experience arises due to gene expression changes across neuronal populations or in a circumscribed neuronal subset is largely unknown. Moreover, it is challenging to establish how changes in the expression of individual genes influence neuronal function and behavior.

Sensory neurons in the periphery are typically tuned to defined stimulus parameters and also exhibit experiencedependent response plasticity. While sensory neurons rapidly adapt to persistent sensory cues via non-transcriptional mechanisms (Martelli and Storace, 2021; Pugh et al., 1999), sensory neurons are now known to also alter their molecular profiles in response to long-term environmental experiences. Circadian light-dark cycles alter phototransduction genes in mammalian photoreceptors; these expression changes may underlie circadian modulation of visual processing (McMahon et al., 2014; Hölter et al., 2012; Storch et al., 2007; Korenbrot and Fernald, 1989). In recent work, it has been shown that the gene expression profiles of olfactory receptor neurons in the mouse olfactory epithelium reflect the animal’s past odor experience, and in turn predict the animal’s response to incoming chemical stimuli (Tsukahara et al., 2021). Whether specific stimulus features are encoded in expression changes of defined sensory gene subsets, and the causal relationships between these changes and behavioral plasticity have yet to be definitively described. It is also currently unclear whether canonical activity-dependent gene regulatory pathways described in central neurons (Flavell and Greenberg, 2008; West et al., 2001) generalize to sensory neurons to modulate their unique response profiles.

*C. elegans* sensory neurons provide an excellent experimental system in which to systematically describe gene expression changes in response to precisely controlled sensory stimuli, and to causally and mechanistically link these expression changes to behavioral plasticity. Sensory neuron types in *C. elegans* are typically comprised of a single left/right neuron pair that are genetically accessible at single neuron resolution (White et al., 1986; Taylor et al., 2021). The sensory responses of many of these neurons have been extensively characterized, and behaviors driven by these neurons are known (Bargmann, 2006; Goodman and Sengupta, 2019). Experience-dependent changes in the expression of a limited number of sensory neuron genes, and the roles of these expression changes in driving behavioral plasticity have been previously described in *C. elegans* (Ryan et al., 2014, Meisel et al., 2014; Gruner et al., 2014; Peckol et al., 2001; Sims et al., 2016; Kaye et al., 2009), suggesting that sensory stimulus-driven activity may globally modulate gene expression programs in individual sensory neuron subtypes to alter behavioral outputs in this organism.

Temperature plays a particularly critical role in regulating both physiology and behavior of *C. elegans* (Hedgecock and Russell, 1975; Ailion and Thomas, 2000; Klass, 1977; Golden and Riddle, 1984a). These nematodes are responsive to multiple aspects of their temperature experience including their actual growth temperature as well as temperature changes. Growth temperature regulates lifespan and a developmental decision between alternate larval life-cycles (Klass, 1977; Ailion and Thomas, 2000; Golden and Riddle, 1984a), and adult animals exhibit robust experiencedependent navigation behaviors in response to temperature changes on spatial thermal gradients (Hedgecock and Russell, 1975). Temperature information is largely transduced by the single AFD thermosensory neuron pair to modulate both development and behavior (Goodman and Sengupta, 2019). We and others previously showed that past and current temperature experience alters the expression of a small number of genes in AFD (Yu et al., 2014; Chen et al., 2016; Kobayashi et al., 2016; Servello et al., 2022), raising the possibility that encoding of different features of the temperature stimulus in the gene expression profile of AFD may underlie the ability of AFD to modulate diverse aspects of the organism’s physiology.

Here we use single neuron type profiling to show that the gene expression program in AFD can reflect multiple aspects of the animal’s temperature experience including the duration, as well as the magnitude of temperature change and its absolute value. We show that the gene expression programs in AFD can encode the absolute growth temperature as well as the magnitude and duration of experienced temperature change, and we causally link expression changes in specific genes to plasticity in AFD-driven functions. We find that expression levels of the conserved DAC-1 Dachsund/DACH1 transcription factor report absolute warm growth temperatures to modulate a larval developmental decision. In contrast, expression changes in an AFD-specific transmembrane protein C12D8.15 (henceforth referred to as PYT-1; see below) report a rapid large magnitude temperature change, and this expression change is essential to drive AFD-mediated thermotaxis behavioral plasticity specifically under these conditions. Temperature alters gene expression in AFD in part via the CRH-1 cAMP response element binding protein (CREB), and we demonstrate that deletion of a single cAMP response element (CRE) motif in the endogenous *pyt-1* promoter is sufficient to abolish temperature-dependent expression changes and neuronal plasticity. Together, these findings describe the mechanistic trajectory from a precisely defined sensory stimulus to neuronal gene expression plasticity, and establish the consequences of these expression changes on organismal and behavioral adaptation at single gene and neuron resolution *in vivo*.

## Results

### Identification of temperature experience-regulated genes in AFD via TRAP-Seq

To assess the temperature-regulated gene expression program in AFD, we identified AFD-enriched mRNAs in animals grown at different temperatures using translating ribosome affinity purification (TRAP) (Heiman et al., 2008; Gracida and Calarco, 2017). TRAP permits flash freezing of animals (see Methods) allowing for more precise measurements of temperature-regulated expression changes as compared to experimental approaches such as cell sorting that require exposure to different temperatures for prolonged time periods (Taylor et al., 2021; Kaletsky et al., 2016). We generated transgenic animals stably expressing the GFP-tagged RPL-1A large ribosomal subunit exclusively in AFD (Figure 1A), and deep sequenced isolated ribosome-associated mRNAs from AFD in adult animals, as well as RNA obtained from whole animal lysates, from cultures grown overnight (∼16 hrs) at either cold (15°C) or warm (25°C) temperatures (Figure 1A; see Methods).

**Fig. 1.**
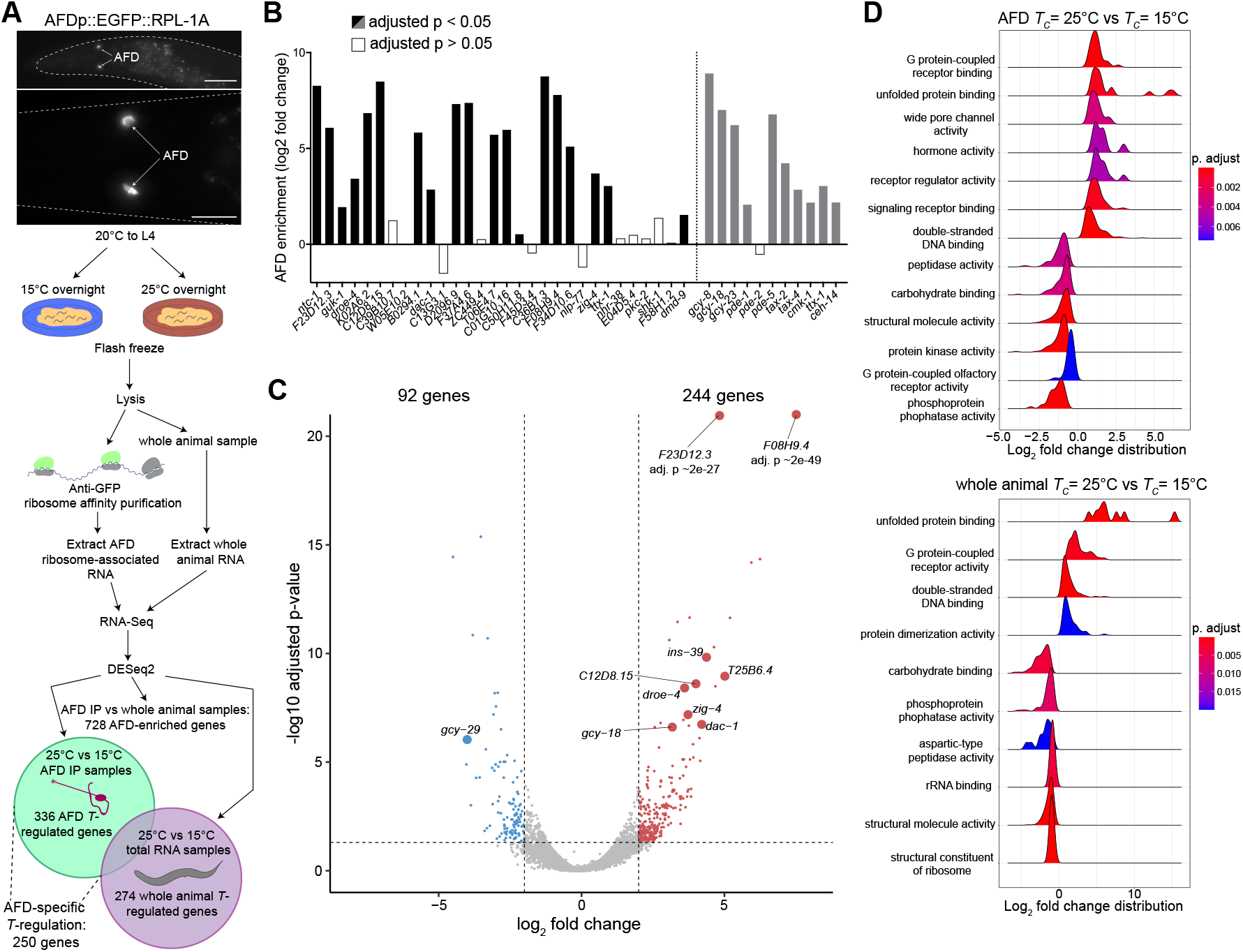
TRAP-Seq profiling of AFD identifies temperature experience-regulated genes. A) (Top) Representative images of the EGFP-tagged RPL-1A ribosomal subunit expressed in AFD in the strain used for TRAP-Seq. (Bottom) TRAP-Seq pipeline to identify temperature-regulated genes in AFD and/or whole animals. IP – immunoprecipation. Scale bars: (top) 50 µm, (bottom) 20 µm; anterior at left. B) Quantification of enrichment in AFD IP vs whole animal samples for the top 30 AFD-enriched genes in the CeNGEN gene expression database (Taylor et al., 2021) (black bars), and for AFD-expressed genes previously implicated in AFD functions (grey bars). Adjusted p-values were calculated by a Wald test performed by DESeq2 (Love et al., 2014). C) Volcano plot of differential gene expression in AFD from animals cultivated at 25°C vs 15°C overnight. Vertical and horizontal dashed lines indicate gene expression changes that are log2 fold change > 2 or < -2 and significant at adjusted p-value < 0.05, respectively. Red and blue dots indicate genes exhibiting significantly higher expression at 25°C or 15°C, respectively. Genes further examined in this work are shown with larger dot sizes. D) Gene set enrichment analysis of temperature-regulated genes in AFD (top) and whole animals (bottom). gseaGO from the clusterProfiler R package (Yu et al., 2012) was used to find enriched gene sets associated with “molecular function” ontology terms. Gene sets with redundant ontology terms were removed manually. Also see Figure S1.

Principal component analysis indicated that the mRNA profiles largely clustered based on tissue (AFD vs whole animal) and temperature experience (15°C vs 25°C) (Figure S1A). Comparison of gene expression in AFD versus whole animal samples identified 728 AFD-enriched genes independent of temperature regulation (Figure 1A). Comparing the set of AFD-enriched genes detected by our TRAP experiments with those identified by the CeNGEN *C. elegans* single gene expression resource (Taylor et al., 2021) showed substantial overlap (Figure 1B). Additionally, previously identified AFD-expressed genes required for AFD development and/or function were also significantly enriched in the AFD-specific TRAP dataset (Figure 1B). Together, these results indicate that we successfully identified AFD-specific genes via TRAP.

We next compared the gene expression profiles of AFD in animals grown at cold and warm temperatures. The expression of a total of 336 genes was altered by temperature experience in AFD, with 250 of these genes exhibiting temperature-regulated expression changes that are likely to be AFD-specific (Figure 1A; Supplementary Data). Of the set of 336 genes, mRNA levels of 244 and 92 genes were present at higher levels in animals grown at warm versus cold temperatures, respectively (Figure 1C). Gene Set Enrichment Analysis (Yu et al., 2012) indicated that temperature experience altered expression of genes implicated in stress response pathways in both AFD and whole animal samples, but specifically altered genes predicted to affect neuronal signaling pathways in AFD (Figure 1D, Figure S1B). Moreover, while neuropeptide and guanylyl cyclase gene classes, including the previously characterized AFD-specific temperature-regulated *gcy-18* thermoreceptor molecule (Yu et al., 2014; Takeishi et al., 2016; Inada et al., 2006), were among genes differentially expressed in AFD (Figure S1B,C), no neuronal gene class was enriched among differentially expressed genes in whole animal samples (Figure S1C). Together, these observations indicate that the expression of a subset of genes implicated in neuronal functions is bidirectionally regulated by temperature experience in the AFD thermosensory neurons.

### Temperature regulates gene expression in AFD on multiple timescales

To validate and further assess temperature-regulated gene expression patterns in AFD, we examined the expression of a subset of identified temperature-regulated genes using fluorescent reporter-tagged genes. Selected genes are predicted to encode proteins that have already been shown, or are hypothesized, to be involved in regulating AFD thermosensory functions. These include genes encoding receptor guanylyl cyclases (*gcy-18, gcy-29*) (Inada et al., 2006; Takeishi et al., 2016), the ortholog of the human DACH1/2 Dachsund transcription factor (*dac-1*) (Colosimo et al., 2004), an insulin-like neuropeptide (*ins-39*) (Servello et al., 2022), a crystallin chaperone protein homologue (*F08H9*.*4*), an immunoglobulin domain protein (*zig-4*) (Aurelio et al., 2002), a predicted novel transmembrane protein (*pyt-1* – PY domain transmembrane protein, see below), and uncharacterized proteins (*F23D12*.*3, T25B6*.*4*, and *droe-4*). Genes were tagged with reporter sequences at their endogenous loci via gene editing or expressed from multicopy extrachromosomal arrays as transcriptional reporter gene fusions (Table S1; Figure S2A). All five endogenously tagged genes, and three of five transcriptional reporter gene fusions exhibited expression exclusively in AFD or in a subset of additional cell types at one or both temperatures (Figure 2A, Figure S2B,C, Table S1). Reporter-tagged proteins were localized to the AFD soma/ nuclei, at the AFD sensory endings, or throughout the cytoplasm (Figure 2A). All examined endogenously tagged genes showed increased expression in AFD upon overnight temperature upshifts used for TRAP-Seq experiments with the exception of *pyt-1::gfp(oy169)* (henceforth referred to as *pyt-1::gfp)* (Figure 2A,B). *dac-1::gfp(oy172)* (henceforth referred to as *dac-1::gfp*) is expressed in AFD as well as in a hypodermal cell and an unidentified neuron type (Figure 2A). Levels of this protein were also increased in these cells upon overnight temperature upshift but to a more modest extent (Figure S2D), indicating that temperature-dependent upregulation of this gene is not restricted to AFD under these conditions.

**Fig. 2.**
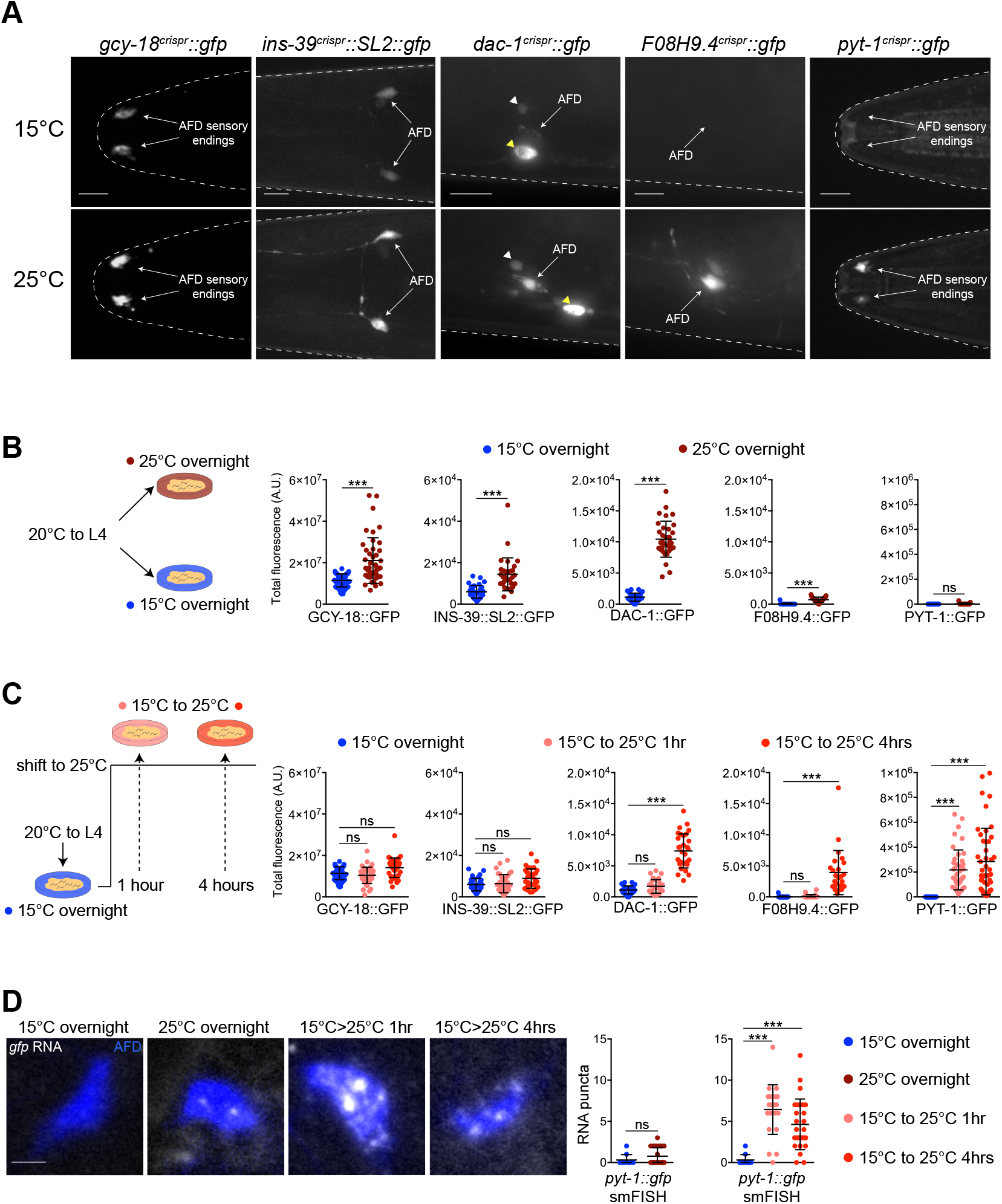
Gene expression in AFD is regulated by the duration of temperature experience. A) Representative images showing expression of indicated endogenously GFP-tagged fusion proteins in the heads of adult hermaphrodites. Animals were grown until the L4 stage at 20°C, shifted to 15°C overnight, and then shifted to 25°C either overnight [*ins-39::SL2::gfp(oy167), gcy-18::gfp(oy165)*] or for 4 hrs [*F08H9*.*4::gfp(syb5551), pyt-1::gfp(oy169), dac-1::gfp(oy172)*]. AFD soma and sensory endings are indicated. *dac-1::gfp* expression in a hypodermal cell and neuron is indicated by a yellow and white arrowhead, respectively. The worm head is outlined with a dotted line. Anterior is at left. Scale bars: 10 µm. B,C) (Left) Cartoons showing temperature exposure protocols. (Right) Quantification of GFP levels of shown fusion proteins in adult animals grown at the indicated conditions. Each dot represents a measurement from a single AFD neuron. Horizontal and vertical lines indicate mean and SD, respectively. n = 22-48 neurons from at least two biologically independent experiments. For each reporter, measurements for conditions shown in B and C were performed together in one imaging session, and the 15°C overnight control data are repeated in these panels. *** indicates different at p<0.001 (one-way ANOVA with Dunnett’s multiple comparisons correction – GCY-18::GFP, INS-39::SL2::GFP, and DAC-1::GFP, or Kruskal-Wallis test with Dunn’s multiple comparisons correction – F08H9.4::GFP and PYT-1::GFP); ns – not significant. D) (Left) Representative images showing smFISH labeling of *pyt-1::gfp* RNA molecules in AFD neurons of L4 animals. Animals were grown under conditions shown in B and C. Scale bar: 2 µm. (Right) Quantification of *pyt-1::gfp* RNA puncta number in AFD. Horizontal and vertical lines indicate mean and SD, respectively. n = 10-27 neurons. *** indicates different at p<0.001 (Kruskal-Wallis test with Dunn’s multiple comparisons correction); ns – not significant. Also see Figure S2.

Experience and/or neuronal activity-dependent gene expression changes occur in distinct and partly overlapping temporal waves (Tyssowski et al., 2018; Fowler et al., 2011). To determine whether AFD-expressed genes display distinct underlying temporal expression dynamics, we next examined the expression of reporter-tagged genes in young adult animals after shifting between cold and warm temperatures for different periods of time. Expression patterns could be categorized broadly into three gene groups based on their temporal patterns of expression changes in AFD following the temperature upshift. The first group included *gcy-18* and *ins-39* whose expression levels in AFD were unaltered even upon a 4 hr shift from 15°C to 25°C (Figure 2C). The second group included *dac-1* and *F08H9*.*4* which exhibited increased expression in AFD at 4 hr after the temperature upshift but not at earlier time points (Figure 2C). We noted that in contrast to the increased expression of DAC-1::GFP in other cell types observed upon overnight exposure to 25°C, levels of this protein were not altered in these cell types at either 1 hr or 4 hr following temperature upshift (Figure S2D). The third group consisted of *pyt-1* whose expression was significantly increased following just 1 hr exposure to 25°C; levels were not further increased with a 4 hr incubation at the warmer temperature (Figure 2C). We infer that the expression levels of a subset of genes in AFD appear to be regulated by the duration of exposure to a new temperature, and that temperature experience regulates gene expression in AFD on multiple timescales and likely via multiple mechanisms.

We next tested whether the observed expression changes arise from transcriptional and/or post-transcriptional changes. We previously showed that upregulation of *gcy-18* expression upon a temperature upshift is primarily mediated via transcriptional mechanisms (Yu et al., 2014). Using smFISH to detect RNAs in AFD, we found that *pyt-1::gfp* RNA was undetectable after cultivation at 15°C in nearly all animals but was strongly upregulated after a 1 or 4 hr shift to 25°C (Figure 2D) recapitulating the expression changes exhibited by the fusion protein. After overnight cultivation at 25°C, *pyt-1::gfp* RNA expression was low but detectable in a subset of animals, although these levels were not significantly different from those in animals cultivated at 15°C (Figure 2D). *pyt-1* may have been identified as an upregulated gene in AFD via TRAP-Seq due to inflation of the fold-change as a consequence of near complete absence of *pyt-1* expression upon cultivation at 15°C. We conclude that warm temperature experience upregulates the expression of a subset of genes in AFD primarily via transcriptional mechanisms.

### Gene expression changes in AFD can reflect the magnitude of experienced temperature change and absolute growth temperature

Gene expression changes across neuronal populations can encode specific features of inducing stimuli, including their magnitude and duration (Tyssowski et al., 2018; Beagan et al., 2020; Fields et al., 1997). Since the AFD neuron pair regulates behavioral and physiological outputs in response to both absolute growth temperature and temperature change (Ailion and Thomas, 2000; Clark et al., 2006; Yu et al., 2014; Servello et al., 2022), we tested whether gene expression programs in this single neuron type reflect multiple features of the temperature stimulus. We focused on *dac-1* and *pyt-1* since each exhibited large expression changes but with markedly distinct temporal dynamics after an experienced temperature shift (Figure 2B,C). To first address whether changes in expression of these genes reflect the magnitude of the temperature change, we shifted animals to a new temperature that was 5°C or 10°C warmer than their initial cultivation temperature for 4 hrs (Figure 3A). We found that DAC-1::GFP levels in AFD increased in proportion to the magnitude of the temperature change (Figure 3A). In contrast, levels of PYT-1::GFP were unaltered with a 5°C temperature shift but showed robust upregulation upon a 10°C temperature shift (Figure 3A). We infer that *dac-1* expression is proportional to the magnitude of temperature change, whereas *pyt-1* is instead only upregulated above a large magnitude temperature change threshold, suggesting analog and digital modes of regulation, respectively.

**Fig. 3.**
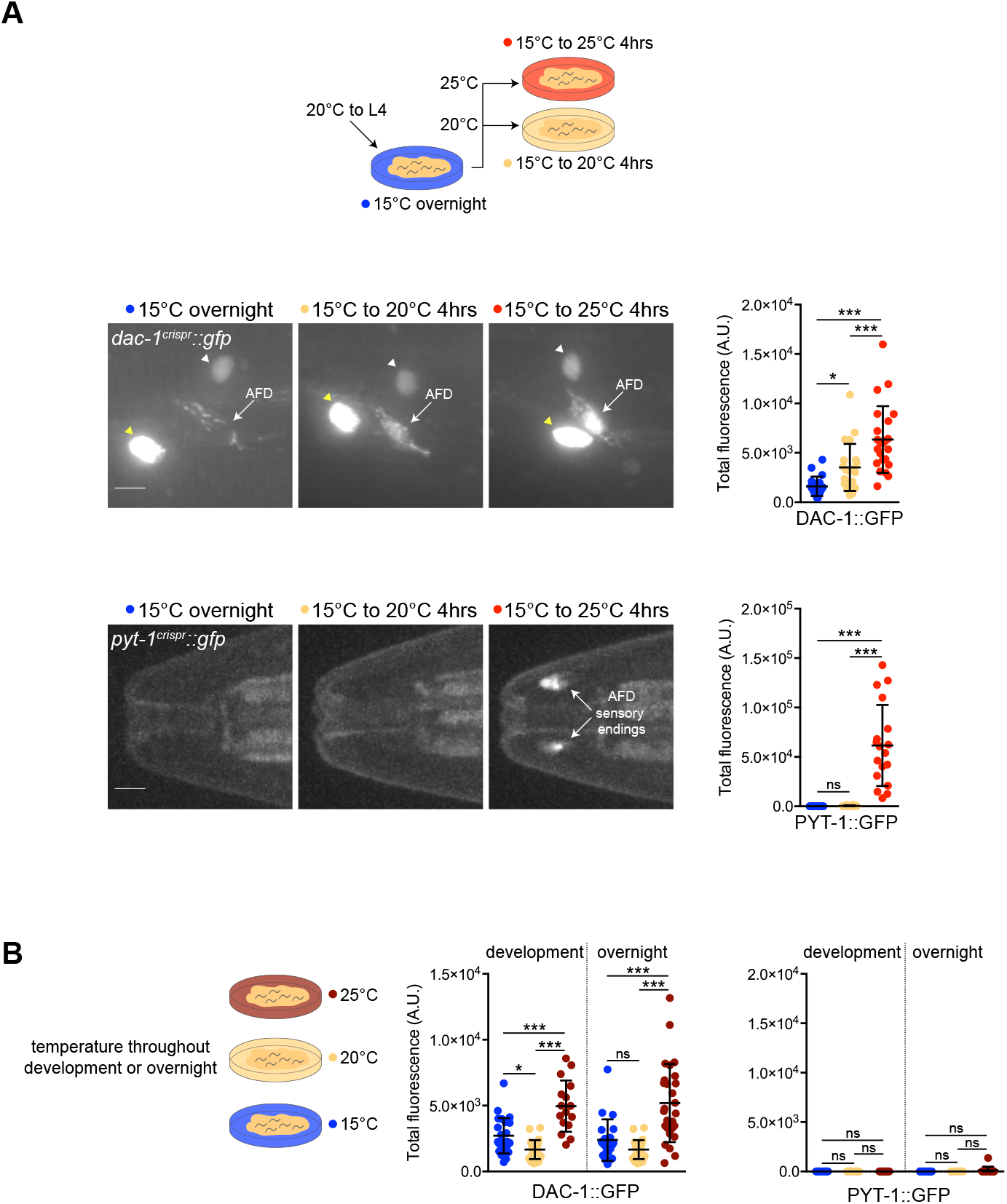
The absolute growth temperature and extent of experienced temperature change are differentially encoded in gene expression patterns in AFD. A) (Top) Cartoon showing temperature exposure protocol. (Middle and bottom) Representative images and quantification of DAC-1::GFP (middle) and PYT-1::GFP (bottom) levels in AFD neurons of adult animals grown at the shown conditions. Yellow and white arrowheads indicate hypodermal and an unidentified neuron expressing DAC-1::GFP, respectively. Scale bars: 10 µm; anterior is at left. Each dot represents a measurement from a single AFD neuron. n = 12-23 neurons from at least two biologically independent experiments. B) (Left) Cartoon showing temperature exposure protocol. (Middle and right) Quantification of DAC-1::GFP and PYT-1::GFP levels in AFD in adult animals grown at the shown conditions. Each dot represents a measurement from a single AFD neuron. Horizontal and vertical lines indicate mean and SD, respectively. n = 10-27 neurons from at least two biologically independent experiments. 20°C measurements are repeated in each reporter expression plot. For all panels, * and *** indicate different at p<0.05 and 0.001, respectively (one-way ANOVA with Dunnett’s multiple comparisons correction – DAC-1::GFP, or Kruskal-Wallis test with Dunn’s multiple comparisons correction – PYT-1::GFP); ns – not significant.

To next test whether levels of these genes can also report the absolute growth temperature as opposed to temperature change, we grew animals at a specific temperature overnight or throughout postembryonic development. DAC-1::GFP levels were low in animals grown under either condition at either 15°C and 20°C, but were consistently elevated in animals grown at 25°C (Figure 3B). Thus, in adult animals that have experienced no change in temperature for a prolonged period of >16 hrs or since hatching, DAC-1 levels are nevertheless elevated in animals cultivated at warm temperatures. In contrast to DAC-1, PYT-1::GFP levels were undetectable in animals grown overnight or throughout postembryonic development at any examined temperature (Figure 3B). We conclude that while levels of DAC-1 can encode both relative temperature change on shorter timescales as well as absolute warm temperature upon prolonged exposure, PYT-1 expression levels only reflect high magnitude relative temperature changes. These observations suggest that different features of the temperature stimulus can be encoded in the gene expression program in AFD, and may enable diverse functional adaptations of AFD to experienced temperature stimuli.

### Temperature experience differentially regulates gene expression in AFD via calcium signaling pathways and a CREB transcription factor

How are different temperature stimulus characteristics transduced to drive expression changes in AFD? Neuronal depolarization and calcium influx modulate gene expression via signaling pathways that include calcium/calmodulindependent protein kinases (West et al., 2001). Similarly, in AFD, we and others previously showed that the temperature-dependent expression changes of a subset of genes are regulated by calcium influx via the TAX-2/TAX-4 cGMP-gated channels and the CMK-1 calcium/calmodulin-dependent protein kinase 1 (Figure 4A) (Satterlee et al., 2004; Yu et al., 2014; Kobayashi et al., 2016; Servello et al., 2022). We tested the extent to which these molecules are also required for translating different aspects of the temperature stimulus into specific gene expression programs in AFD.

**Fig. 4.**
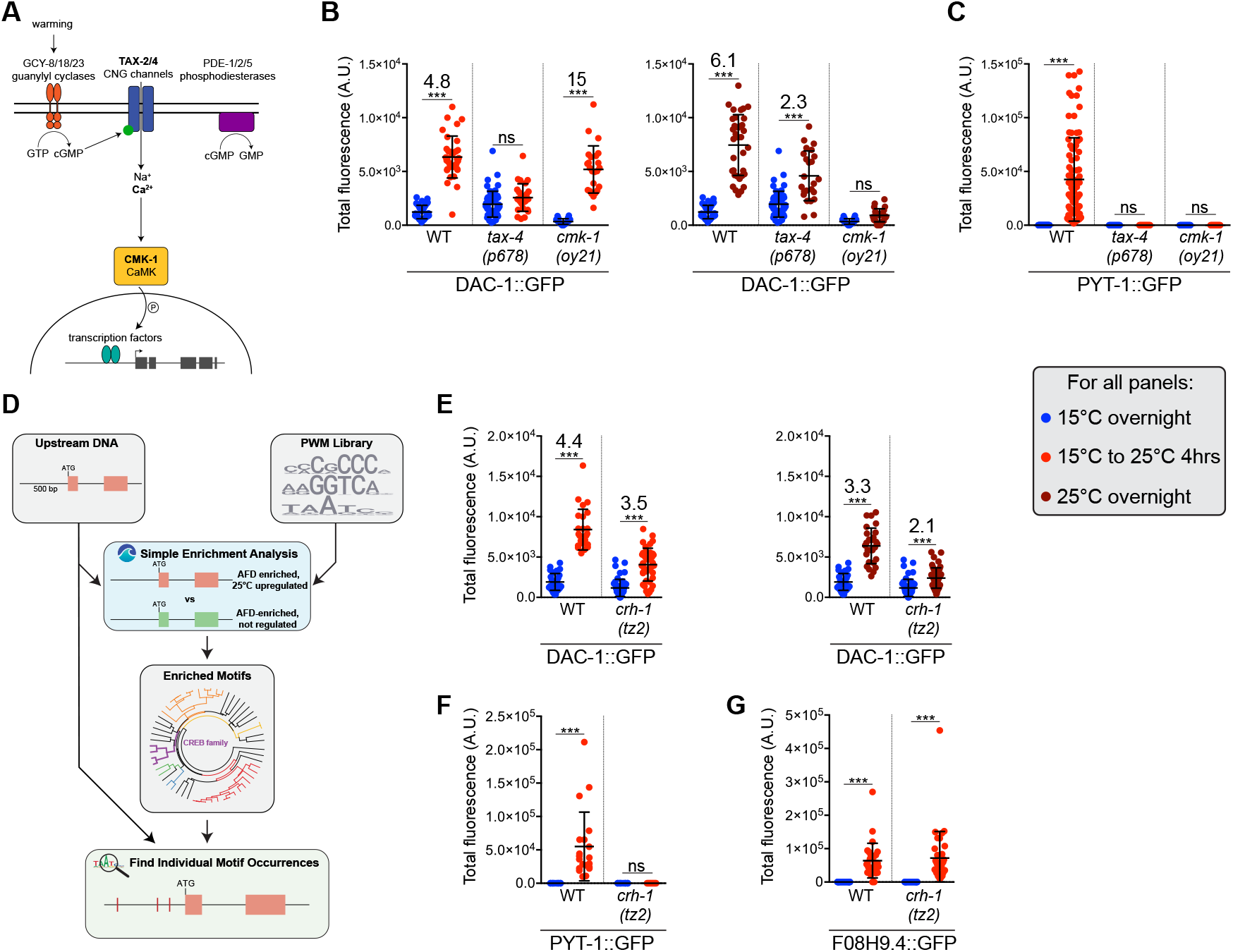
Temperature stimulus features are transduced differentially by calcium signaling pathways and CREB to regulate gene expression. A) Cartoon of the AFD thermosensory signal transduction pathway. B,C,E,F) Quantification of DAC-1::GFP (B,E) and PYT-1::GFP (C,F) levels in adult animals grown at the indicated conditions in tax-4 and cmk-1 (B,C), and crh-1 (E,F) mutants. All reporters were tagged endogenously with GFP. Each dot is a measurement from a single AFD neuron. Horizontal and vertical lines indicate mean and SD, respectively. n = 9-85 neurons from at least two biologically independent experiments. A subset of WT data for DAC-1::GFP measurements is repeated from Figure 2B,C. *** indicates different at p<0.001 (one-way ANOVA with Dunnett’s multiple comparisons correction – DAC-1::GFP, Mann-Whitney test – PYT-1::GFP); ns – not significant. Numbers at top indicate fold-change between indicated conditions. D) Schematic of the analysis pipeline used to predict transcription factor binding sites upstream of temperature-regulated genes (see Methods). PWM: position weight matrix. G) Quantification of endogenously tagged F08H9.4::GFP levels in adult animals grown at the indicated conditions in *crh-1* mutants. Each dot is a measurement from a single AFD neuron. Horizontal and vertical lines indicate mean and SD, respectively. n = 20-34 neurons from at least two biologically independent experiments. *** indicates different at p<0.001 (Mann-Whitney test).

Consistent with a requirement for calcium influx upon a temperature change to alter gene expression, upregulation of both PYT-1::GFP as well as DAC-1::GFP upon a temperature upshift was fully dependent on TAX-4 (Figure 4B,C). We also previously showed that elevated expression of *ins-39* upon a temperature shift requires TAX-4 (Servello et al., 2022). However, the requirement for CMK-1 was distinct such that only PYT-1::GFP but not DAC-1::GFP expression upon a temperature upshift required CMK-1 (Figure 4B,C). In contrast, a mutation in tax-4 only partially disrupted elevation of DAC-1::GFP levels upon prolonged growth at warm temperatures, although this upregulation was fully dependent on CMK-1 (Figure 4B,C). We conclude that calcium influx through TAX-4 acts via both CMK-1-dependent and -independent pathways to alter gene expression upon a temperature upshift, but that absolute warm temperature information is encoded via activation of CMK-1 by alternate pathways (see Figure S3D for summary).

We next investigated the downstream transcriptional regulatory mechanisms that drive temperature experiencedependent gene expression changes. Restricting our search space to sequences 500 bp upstream of the translation start site of each temperature-upregulated gene in AFD, we first identified matches to a curated list of transcription factor binding motifs identified from analyses in multiple organisms (Figure 4D) (Bailey and Grant, 2021; Weirauch et al., 2014). 57 motifs were specifically enriched in the set of AFD-expressed and temperature-regulated but not temperature-insensitive genes (Figure 4D, see Methods). These motifs were then clustered into 27 groups representative of transcription factor families that bind a similar motif (Figure 4D) (Castro-Mondragon, 2017). Of the motif clusters identified, we noted one set that is predicted to be recognized by transcription factors related to the neuronal activity-dependent transcription factor cyclic AMP response element-binding protein (CREB) (Sheng et al., 1990; Kaang et al., 1993) (Figure 4D). The C. elegans CREB ortholog CRH-1 is expressed in AFD and has previously been implicated in regulating thermosensory behaviors and temperature-and AFD-dependent developmental pathways (Nishida et al., 2011; Nakano et al., 2022; Chen et al., 2016; Park et al., 2021).

We tested whether CRH-1 mediates temperature experience-dependent regulation of a subset of identified AFD-expressed genes. In response to a temperature upshift for 4 hrs, elevated expression of *dac-1* and *pyt-1* was partly or fully CRH-1-dependent, respectively (Figure 4E,F). However, at this timepoint, increased expression of *F08H9*.*4* was CRH-1-independent (Figure 4G), indicating that factors other than CRH-1 also read out temperature shift information to alter gene expression at this timepoint. CRH-1 also partly abrogated increased DAC-1::GFP levels upon overnight growth at warm temperatures indicating that in addition to mediating temperature change information (Figure 4E), CRH-1 may also integrate warm growth temperature information to regulate gene expression. These results highlight the diversity of regulatory pathways that translate specific aspects of the temperature stimulus into unique gene expression programs in AFD (summarized in Figure S3D).

### CREB acts both directly and indirectly to regulate gene expression in response to different features of the temperature stimulus

We next tested whether CRH-1 acts directly or indirectly to regulate temperature experience-dependent expression of *dac-1* and *pyt-1* in AFD. The *dac-1* locus is predicted to encode two isoforms (DAC-1A and DAC-1B) that differ at their N-termini (www.wormbase.org). A transcriptional reporter driven under 2.4 kb of regulatory sequences upstream of the dac-1a isoform drove expression only in hypodermal cells, whereas a reporter driven under 450 bp of promoter sequences that include part of the dac-1b 5’UTR and upstream DNA drove expression specifically in AFD (Figure S3A). Moreover, expression of the dac-1bp::gfp reporter was strongly upregulated following a temperature upshift (Figure S3B). Deleting 4 of 5 predicted CREB binding sites (CREs - cAMP response elements; Figure 5A,B) at the endogenous dac-1b promoter locus resulted in significant loss of temperature-regulation of DAC-1::GFP after a 4 hr temperature upshift (Figure 5C). A similar result was obtained upon mutating only two of the highest confidence predicted CREs at the endogenous locus (Figure 5C). However, these CRE mutations had little to no effect on *dac-1::gfp* expression levels in response to absolute warm growth temperatures (Figure 5C). We also identified one high confidence CRE and four additional lower confidence CREs in the upstream 500 bp regulatory sequences of *pyt-1* (Figure 5D). Mutating the high confidence CRE at the endogenous locus resulted in nearly complete loss of temperature-dependent upregulation of *pyt-1* (Figure 5E). This CRE was also conserved in the upstream sequences of *pyt-1* orthologs in a subset of nematode species (Figure S3C). We conclude that CRH-1 directly binds to CREs to regulate both *dac-1* and *pyt-1* expression upon a temperature shift, but that CRH-1 may act indirectly to modulate *dac-1* levels in response to warm growth temperature.

**Fig. 5.**
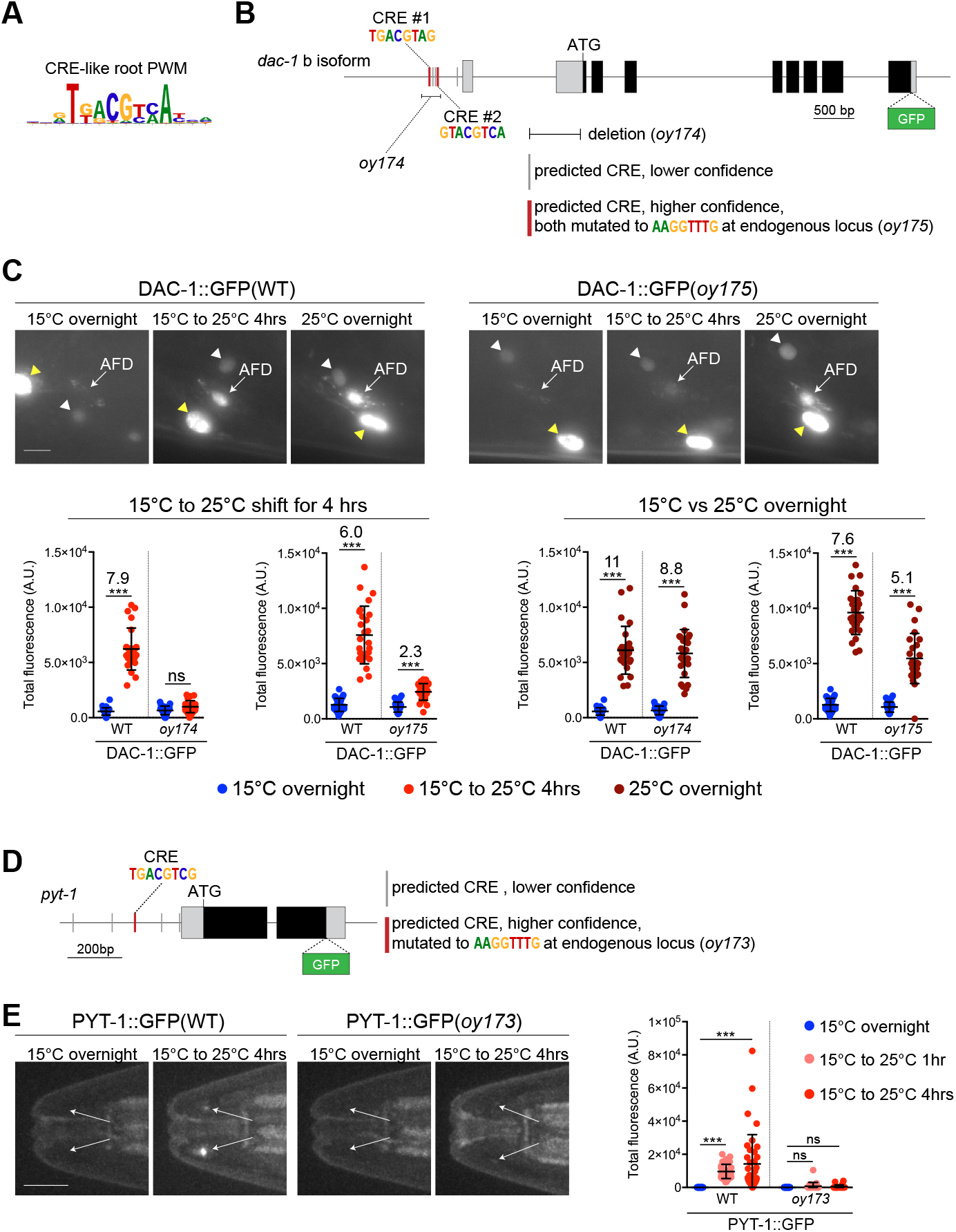
CREB can act directly to regulate gene expression in response to a temperature upshift. A) Root PWM derived from CRE-related PWMs generated with RSAT (Castro-Mondragon et al., 2017). B) Genomic structure of the dac-1b::gfp locus. Predicted CRE motifs within 450 bp upstream of the first exon of *dac1b* are depicted as vertical bars. The extent of the oy174 deletion and mutations in two predicted CREs (oy175) indicated by thick vertical red bars were generated in the endogenous *dac-1::gfp(oy172)* reporter (see Methods). Thin lines, grey boxes, and black boxes indicate upstream sequences/introns, untranslated regions, and coding sequences, respectively. C) (Left) Representative images of endogenously tagged DAC-1::GFP expression under wildtype or mutated (2XCRE mutant; oy175) regulatory sequences in adult animals grown at the indicated conditions. White and yellow arrowheads indicate a hypodermal cell and neuron, respectively. Scale bar: 5 µm. Anterior is at left. (Right) Quantification of endogenously tagged DAC-1::GFP levels in AFD in adult animals expressed under wild-type, or mutated (deleted CREs - oy174; two mutated CREs - oy175). Each dot is a measurement from a single AFD neuron. Horizontal and vertical lines indicate mean and SD, respectively. n = 23-27 neurons from at least two biologically independent experiments. A subset of WT data for DAC-1::GFP measurements is repeated from Figure 2B,C. *** indicates different at p<0.05 and 0.001, respectively (one-way ANOVA with Dunnett’s multiple comparisons correction). ns – not significant. Numbers at top indicate fold-change between indicated conditions. D) Structure of the *pyt-1::gfp* gene locus. Predicted CREs are depicted as vertical bars. The DNA sequence indicated by the thick vertical red bar was mutated in the endogenous *pyt-1::gfp* reporter to generate oy173 (see Methods). Thin lines, grey boxes, and black boxes indicate upstream sequences/introns, untranslated regions, and coding sequences, respectively. E) (Left) Representative images of endogenously tagged PYT-1::GFP expression at the AFD sensory endings (arrows) under wild-type or mutated (CRE mutant; oy173) regulatory sequences in adult animals grown at the indicated conditions. Scale bar: 10 µm. Anterior is at left. (Right) Quantification of endogenously tagged PYT-1::GFP levels expressed under wild-type or mutated (CRE mutant; oy173) regulatory sequences in adult animals grown at the indicated conditions. Each dot is a measurement from a single AFD neuron (arrows). Horizontal and vertical lines indicate mean and SD, respectively. n = 20-34 neurons from at least two biologically independent experiments. *** indicates different at p<0.001 between indicated conditions; ns – not significant (Kruskal-Wallis test with Dunn’s multiple comparisons correction); ns – not significant. Also see Figure S3.

### Warm growth temperature information is integrated in DAC-1 expression levels in AFD to modulate a developmental decision

The differential regulation of genes such as *dac-1* and *pyt-1* in AFD by different temperature stimulus features suggests that these molecules may regulate distinct AFD properties as a function of the animal’s temperature experience. Since DAC-1 levels reflect both a short-term temperature shift as well as absolute warm growth temperatures, we explored the contributions of this protein to shaping AFD-driven processes.

When placed on thermal gradients. C. elegans preferentially moves towards the temperature they experienced in the past 3-4 hrs; this behavioral preference is altered upon exposure to a new temperature for a similar time period (Hedgecock and Russell, 1975). To test whether increased expression of *dac-1* upon a temperature upshift for 4 hrs influences thermotaxis behavioral plasticity, we examined the navigation behavior of *dac-1* animals grown at 15°C and shifted to 25°C for 4 hrs in high resolution thermotaxis behavioral assays. However, *dac-1* mutants exhibited no defects in thermotaxis behavior upon growth at either 15°C or following a temperature upshift to 25°C for 4 hrs (Figure S4A). DAC-1 may regulate AFD-driven behaviors under specific conditions or mediate as yet unknown AFD functions in adult animals experiencing this temperature change.

Warmer growth temperatures (25°C) enhance entry of C. elegans larvae into the stress-resistant dauer developmental stage under defined environmental conditions (Golden and Riddle, 1984a; Golden and Riddle, 1984b; Fielenbach and Antebi, 2008). Thus, growing larvae at 27°C, or at 25°C with added exogenous pheromone preferentially promotes dauer arrest over continued reproductive growth (Ailion and Thomas, 2000; O’Donnell et al., 2018; Golden and Riddle, 1984b; Ailion and Thomas, 2000). As in adult animals, *dac-1* expression was also increased in AFD in L1 larvae grown at warm temperatures (Figure 6A), suggesting that DAC-1 may influence this temperature-regulated developmental decision. We found that a significantly larger fraction of *dac-1* mutant animals entered the dauer stage when grown at 27°C (Figure 6B); this phenotype was similar to those of *ttx-1* mutants in which the AFD neurons exhibit developmental defects (Ailion and Thomas, 2000). The enhanced dauer formation defect of *dac-1* mutants was fully rescued by expression of DAC-1 specifically in AFD (Figure 6B). Similarly, more *dac-1* mutants exhibited dauer entry at 25°C in the presence but not absence of low added concentrations of the ascr5 pheromone (Figure 6C,D). Dauer formation is driven in part via downregulation of TGF-beta and insulin signaling from the ASI and ASJ sensory neurons in the head of C. elegans (Fielenbach and Antebi, 2008; O’Donnell et al., 2018). However, expression of neither *daf-7* TGF-beta nor of the *daf-28* insulin-like peptide gene was significantly altered in *dac-1* L1 larvae (Figure S4B,C), indicating that DAC-1 may regulate other neuroendocrine genes to suppress dauer formation at warm growth temperatures. We infer that DAC-1 levels encode warm growth temperature information in the AFD neurons of larvae to appropriately modulate the dauer developmental decision.

**Fig. 6.**
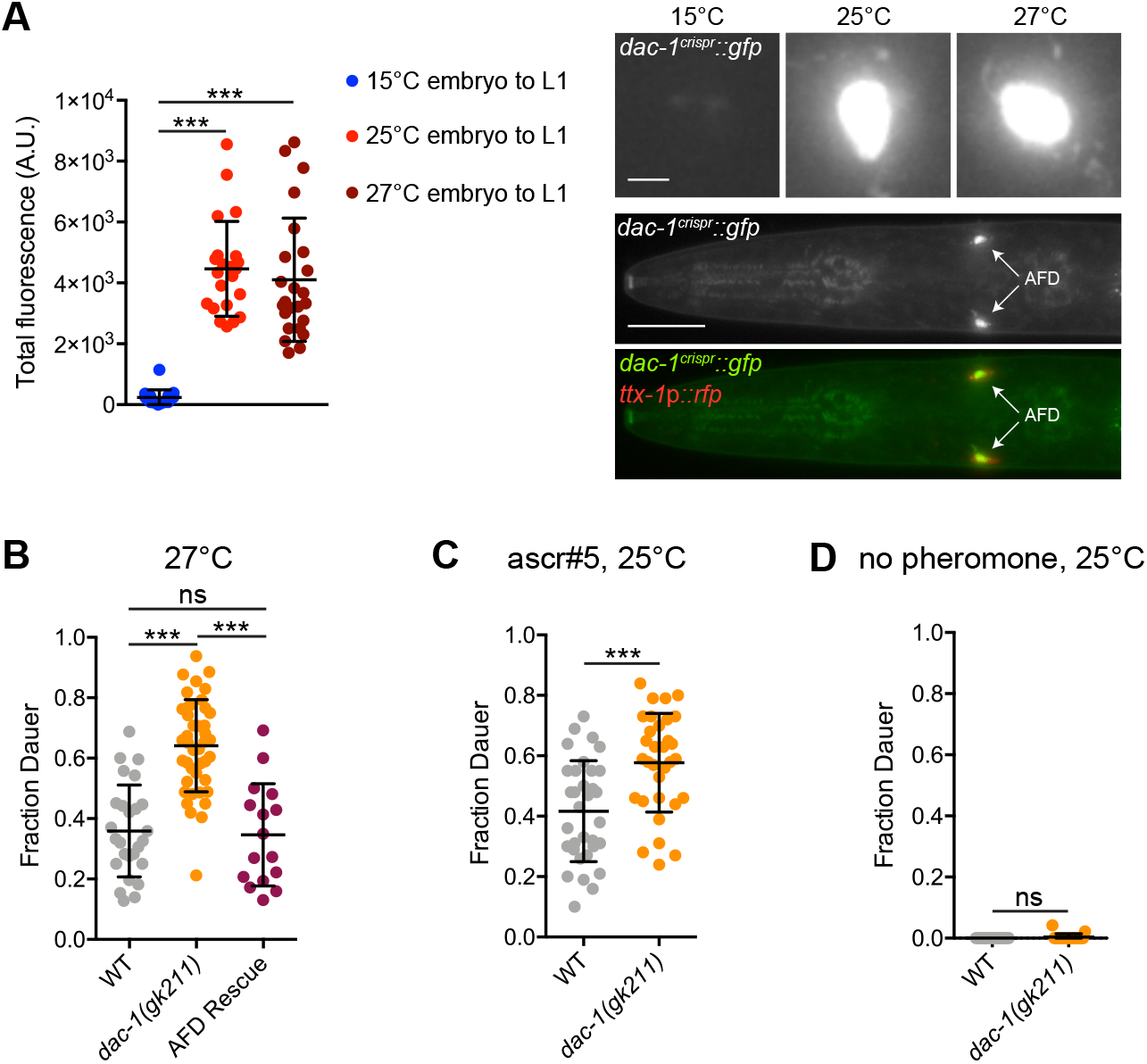
Upregulation of *dac-1* in response to warm growth temperatures regulates the dauer developmental decision. (Left) Quantification of DAC-1::GFP levels in AFD in L1 animals grown at the shown conditions. Each dot represents a measurement from a single AFD neuron. Horizontal and vertical lines indicate mean and SD, respectively. n = 19-24 neurons from at least two biologically independent experiments. (Right top) Representative images of DAC-1::GFP expression in AFD of L1 larvae at the indicated temperatures. (Right bottom) Colocalization of DAC-1:GFP with ttx-1p::rfp in AFD soma. Scale bar: (top) 2 µm, (bottom) 100 µm. B-D) Dauers formed by animals of the indicated genotypes. Each dot indicates the fraction of dauers formed in a single assay. Horizontal and vertical lines indicate mean and SD, respectively. n = 16-42 assays from at least two biologically independent experiments. dac-1a(cDNA)::SL2::mCherry was expressed under the gcy-8 promoter in C. For all panels, *** indicates different at p<0.001, (A - one-way ANOVA with Dunnett’s multiple comparisons correction, B - one-way ANOVA with Tukey’s multiple comparisons correction, C,D - t test); ns – not significant. Also see Figure S4.

### Direct CREB-mediated regulation of *pyt-1* expression in AFD links the experience of a specific temperature change to thermosensory behavioral plasticity

The rapid upregulation of *pyt-1* expression only in response to a large magnitude temperature upshift suggested that this molecule may specifically link transcriptional changes to neuronal and behavioral plasticity in response to experienced temperature change. *pyt-1* encodes a small single transmembrane domain protein (Figure 7A) and is conserved in a subset of both free-living and parasitic nematodes but not in other species (Figure S5A). The predicted C-terminal domain of PYT-1 contains two sequences (PPxY and LPxY, together referred to as a PY motif) (Figure 7A, Figure S5B) which have been shown to bind the conserved Group 1 WW protein-protein interaction motif present in many proteins with a range of functions (Salah et al., 2012; Huang et al., 2020). To begin to explore a function for PYT-1, we first characterized the localization of PYT-1 at the AFD sensory endings. Thermosensory signaling proteins exhibit differential localization within the AFD sensory endings such that the thermosensor receptor guanylyl cyclases including GCY-18 are localized to finger-like microvilli but are excluded from the small cilium, whereas the TAX-2 and TAX-4 thermotransduction channels are restricted to the proximal ciliary region (Figure 7B) (Nguyen et al., 2014). Functional PYT-1::GFP (see Figure 7H) was enriched in a region at the base of the AFD cilium and also present in dimmer puncta distributed throughout the AFD microvilli but was excluded from the cilium itself (Figure 7B,C).

**Fig. 7.**
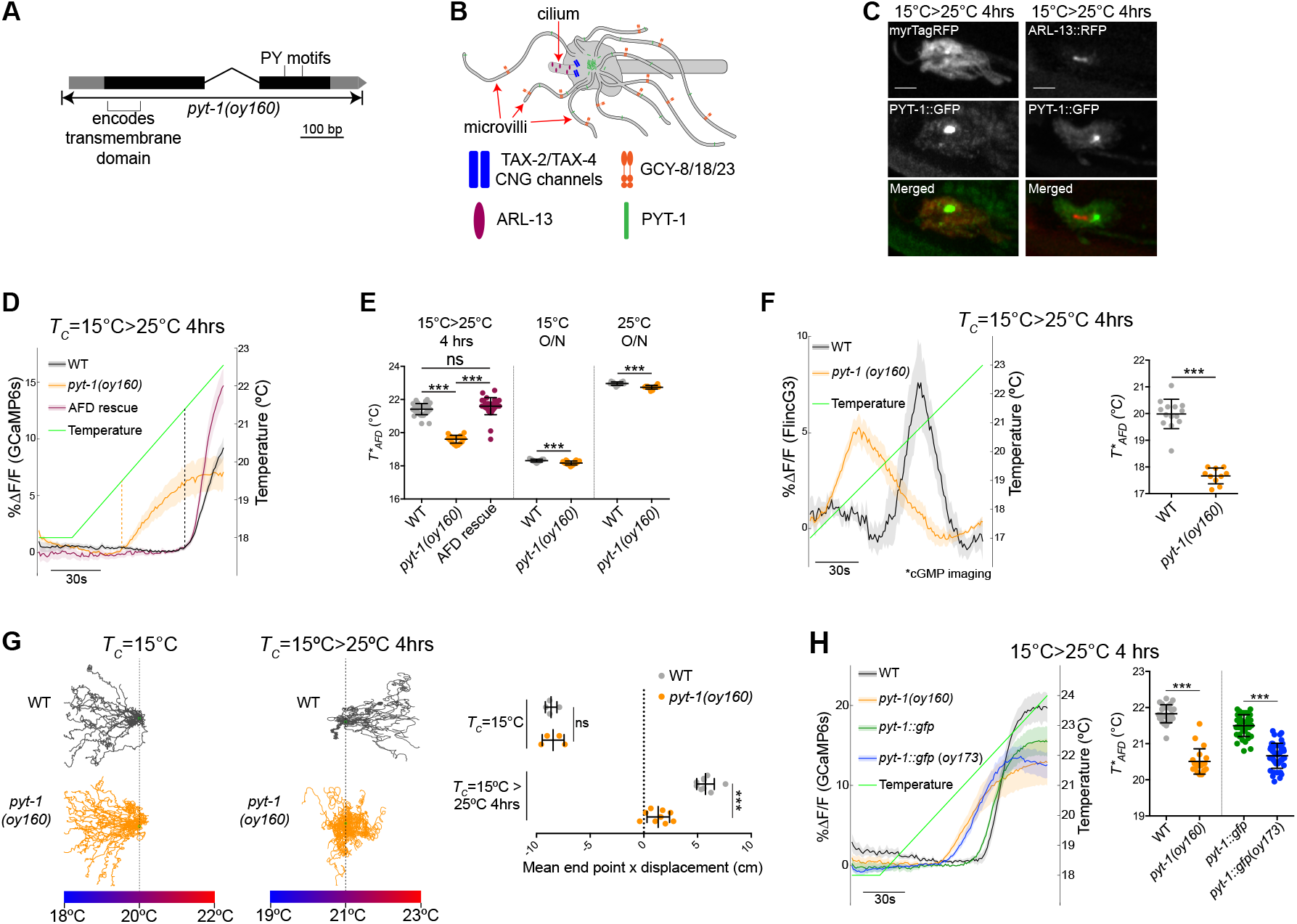
CREB-mediated upregulation of the *pyt-1*-encoded transmembrane protein drives thermosensory behavioral plasticity in response to a specific temperature experience. A) *pyt-1* genomic locus and oy160 deletion boundaries. Black and grey boxes indicate coding DNA and untranslated regions, respectively. Sequences encoding the predicted transmembrane domain and location of the PY motifs are indicated. B) Cartoon of the AFD sensory ending and localization of thermotransduction proteins (Nguyen et al., 2014; this work). C) Representative images of PYT-1::GFP localization within the AFD sensory endings at the indicated temperature conditions. AFD membrane and cilia were visualized with ttx-1p::myrTagRFP and ttx-1p::ARL-13::RFP, respectively. Scale bars: 2 µm. Anterior is at left. D) GCaMP traces acquired from AFD in adult animals grown under the indicated conditions during the shown temperature ramp protocol (green line). Thick lines and shading indicate the average deltaF/F change and SEM, respectively. Dashed vertical lines indicate the corresponding T*AFD for wild-type and *pyt-1(oy160)* animals. The AFD rescue transgene was *gcy-8p::pyt-1(genomic)::SL2::mCherry*. E) Quantification of T*AFD for animals grown under the indicated conditions. Each dot is a measurement from a single animal. Horizontal and vertical lines indicate mean and SD, respectively. n = 12-37 animals from at least two biologically independent experiments. F) (Left) FlincG3 cGMP indicator traces from AFD in adult animals during the shown temperature ramp protocol (green line). Traces are corrected for photobleaching by subtraction of an exponential fit curve for visualization (see Methods). Thick lines and shading indicate the average deltaF/F change and SEM, respectively. (Right) Quantification of T*AFD from traces shown at left. Each dot is a measurement from a single animal. Horizontal and vertical lines indicate mean and SD, respectively. n = 10-14 animals from at least two biologically independent experiments. G) (Left) Representative thermotaxis behavior trajectories of adult wild-type and *pyt-1(oy160)* animals in a single assay on thermal gradients grown at the indicated conditions. Vertical dashed lines indicate the starting temperature on the gradient. (Right) Quantification of thermotaxis assay end points relative to starting position. Each dot indicates the average end point distance of all animals in a single assay. Vertical and horizontal lines indicate mean and SD, respectively. n = 4-9 assays of 15-25 animals each from at least two independent days. H) (Left) GCaMP traces from AFD in adult animals during the temperature ramp protocol (green line). Thick lines and shading indicate the average deltaF/F change and SEM, respectively. (Right) Quantification of T*AFD from traces shown at left. Each dot is a measurement from a single animal. Horizontal and vertical lines indicate mean and SD, respectively. n = 20-44 animals from at least two biologically independent experiments. For all panels, *** indicates different at p<0.001 (E - one-way ANOVA with Tukey’s multiple comparisons correction - 15°C>25°C 4hrs, t test - 15°C O/N and 25°C O/N, F-I - t test,); ns – not significant. Also see Figure S5.

To investigate *pyt-1* function, we generated a null allele of *pyt-1* via gene editing (Figure 7A). The overall morphology of the AFD sensory endings was not significantly altered in *pyt-1* mutants (Figure S5C). We next assessed temperature-driven changes in intracellular calcium dynamics in the AFD sensory endings using the genetically encoded GCaMP6s calcium sensor. In wild type animals, AFD is rapidly activated by warming stimuli above a precise threshold temperature (T*AFD) that is altered in a temperature experience-dependent manner on a fast minutes- and slower timescale of 3-4 hrs (Kimura et al., 2004; Clark et al., 2006; Ramot et al., 2008; Yu et al., 2014). This flexibility in T*AFD together with plasticity in AFD synaptic output drives plasticity in thermotaxis navigation behaviors (Ramot et al., 2008; Hawk et al., 2018; Clark et al., 2006; Kimura et al., 2004; Ohnishi et al., 2011). We found that the T*AFD of *pyt-1* mutants was significantly lower than that of wild-type animals when animals experienced a 15°C to 25°C up-shift for 4 hrs (Figure 7D,E), but showed no defects in the fast component of T*AFD adaptation following a temperature upshift for 10 mins (Figure S6C). Moreover, unlike the rapid activation of calcium responses in wild type animals, AFD neurons in *pyt-1* mutants responded to warming with a gradual increase in calcium (Figure 7D). Response phenotypes were fully rescued upon AFD-specific expression of GFP-tagged PYT-1 under a temperature-insensitive promoter (Figure 7E). Remarkably, T*AFD as well as the dynamics of the calcium response in AFD were only altered to a minor extent in *pyt-1* mutants when grown overnight at either 15°C or 25°C when expression levels of this gene are low (Figure 7E, Figure S6A,B). These observations suggest that PYT-1 plays a specific role in tuning AFD thermosensory response plasticity at an hours-long timepoint after a temperature upshift that coincides temporally with upregulated expression of this gene.

AFD responds to warming stimuli by increasing intracellular cGMP levels via the thermoreceptor guanylyl cyclases; cGMP in turn opens the TAX-2 and TAX-4 cGMP-gated channels to allow calcium influx and depolarization (Goodman and Sengupta, 2019). To determine whether PYT-1 functions upstream or downstream of cGMP production in the signal transduction cascade, we next imaged changes in cGMP levels at the AFD sensory endings in response to a warming stimulus using the genetically encoded FlincG3 fluorescent cGMP sensor (Bhargava et al., 2013; Woldemariam et al., 2019). The AFD neurons in *pyt-1* mutants were again activated at a significantly lower temperature compared to wild type following a 4 hr temperature upshift (Figure 7F). Similar to calcium response dynamics, cGMP levels also rose gradually in AFD in *pyt-1* mutants compared to the rapid increase observed in wild-type AFD (Figure 7F), indicating that PYT-1 likely acts at the first step of AFD thermosensory signal transduction upon a temperature upshift for 4 hrs. PY motif containing proteins can be direct targets of protein degradation by the Nedd4 family of E3 ubiquitin ligases via interaction with their WW domains (Rotin et al., 2001; Harty et al., 2000). A subset of small PY-containing transmembrane proteins that otherwise do not share extensive sequence homology have also been shown to act as adapters to recruit these ligases to target proteins that may themselves lack the PY motif (Myat et al., 2002; Hettema et al., 2004; Gorla et al., 2022). However, the levels of the endogenously tagged GCY-18 guanylyl cyclase was unaltered in *pyt-1* mutant animals after a shift from 15°C to 25°C for 4 hrs (Figure S5C), suggesting that PYT-modulates cGMP levels at the AFD thermosensory endings via alternative mechanisms. Resetting T*AFD as a function of temperature experienced in the past 3-4 hrs is critical for animals to appropriately shift their temperature preference in thermotaxis navigation behaviors (Goodman and Sengupta, 2019). *pyt-1* mutants were markedly defective in their ability to shift their navigation preference towards warmer temperatures (positive thermotaxis) upon a shift from 15°C to 25°C for 4 hrs consistent with defects in their ability to appropriately adapt T*AFD (Figure 7G), but moved similarly to wild-type animals on an isothermal plate at 20°C (Figure S6D). However, reflecting low or absent *pyt-1* expression upon overnight growth at 15°C and no effect on T*AFD in *pyt-1* mutants under these conditions, thermotaxis navigation was unaffected in *pyt-1* animals cultivated at 15°C (negative thermotaxis; Figure 7G).

If temperature-dependent upregulation of *pyt-1* via CREB binding to a single CRE is important for regulating AFD thermosensory plasticity, we reasoned that the *pyt-1* allele in which just the CRE has been mutated (oy173) should exhibit temperature response phenotypes in AFD similar to those of the *pyt-1(oy160)* protein null allele. As shown in Figure 7H, both T*AFD and the dynamics of temperature-evoked intracellular calcium changes in AFD were similarly altered in animals carrying either allele upon a temperature upshift from 15°C to 25°C for 4 hrs. Moreover, similar to *pyt-1* null mutants, AFD temperature responses were largely unaffected in *pyt-1(oy173)* mutants upon overnight growth at either 15°C or 25°C (Figure S6E). crh-1 mutants also exhibited significant defects in T*AFD upon a 15°C to 25°C shift for 4 hrs as well as upon overnight growth at 25°C but a weaker defect upon overnight growth at 15°C (Figure S6F) (Nishida et al., 2011). Together, we conclude that temporally regulated CREB-mediated transcriptional upregulation of *pyt-1* specifically regulates plasticity in AFD thermosensory responses and thermotaxis behavior at a defined timepoint following a large magnitude temperature upshift.

## Discussion

Here we show that specific aspects of an animal’s temperature experience, including the duration and absolute value of the experienced temperature, as well as the extent of temperature change, are quantitatively encoded in the gene expression profile of the single pair of AFD thermosensory neurons in C. elegans. Functional characterization of two genes identified in this work show that their patterns of expression modulation are critical for driving AFD-dependent behavioral and developmental decisions in response to defined aspects of the animal’s temperature experience. We further find that a subset of this experience-dependent regulation is mediated by the CRH-1 CREB transcription factor, and that mutating an individual CREB binding site is sufficient to abolish activity-dependent expression, resulting in defects in neuronal plasticity. Our findings demonstrate encoding of sensory stimulus features in unique gene expression programs thereby coupling stimulus to transcription (Tyssowski and Gray, 2019) in a single sensory neuron type in vivo, and mechanistically describe how these expression changes in single genes influence neuronal and organismal plasticity.

Our results suggest that in AFD, the extent of experienced temperature change is reflected in analog changes in the expression of genes such as *dac-1* thereby providing a mechanism to faithfully report this stimulus feature and modify neuronal function along a continuous axis. Graded CREB-dependent changes in the expression of the *flp-6* neuropeptide gene in AFD at different growth temperatures have previously also been shown to be important for AFD-mediated regulation of longevity at warm temperatures (Chen et al., 2016). Similarly, in rodent olfactory receptor neurons, analog changes in gene expression in response to odor experience history reflect the animal’s odor history and regulate future olfactory responses (Tsukahara et al., 2021). In contrast, a digital regulatory design reports a large magnitude temperature change by upregulating *pyt-1* expression only above a specific stimulus change threshold (Burz et al., 1998; Houchmandzadeh et al., 2002). The magnitude of temperature change has previously been suggested to be proportional to the amplitude of the calcium response in AFD (Clark et al., 2006). Graded changes in second messenger levels could be translated into analog changes in gene expression via non-cooperative binding of transcription factors to multiple cisregulatory elements (Stewart-Ornstein et al., 2013; Giorgetti et al., 2010). *dac-1* levels also report the absolute warm temperature upon prolonged exposure. Resting calcium levels in AFD have recently been reported to be correlated with growth temperature (Thapliyal et al., 2022; Ippolito et al., 2021). Basal calcium levels in AFD at different temperatures may drive differential expression of downstream targets via distinct regulatory molecules including CMK-1. The ability to couple different features of the thermal stimulus to distinct gene expression programs allows AFD to precisely modulate downstream signaling pathways as a function of the animal’s specific temperature experience.

The critical role of specific gene expression changes in AFD in response to defined stimulus features is exemplified in our functional characterization of both *dac-1* and *pyt-1*. Upregulation of *dac-1* expression in larvae upon prolonged growth at warm temperatures allows animals to integrate their long-term temperature experience to appropriately modulate the dauer decision pathway. Conversely, the rapid and transient upregulation of *pyt-1* following a large magnitude temperature upshift for 1-4 hrs is essential to correctly tune plasticity in AFD thermosensory responses and thermotaxis behavior specifically on this timescale. While it has been suggested that adaptation of only the synaptic output of AFD on an hours-long timescale is sufficient to drive plasticity in thermotaxis behaviors upon a temperature upshift (Hawk et al., 2018), our experiments further confirm that adaptation of the sensory response threshold of AFD on this timescale is also critical for behavioral plasticity (Yu et al., 2014). Homeostatic mechanisms can precisely restore neuronal and/or circuit excitability after activity perturbation by tuning gene expression (O’Leary et al., 2014; Kulik et al., 2019; Tsukahara et al., 2021). Adaptation of the thermosensory response threshold, dependent in part on alteration of expression of genes such as *pyt-1*, allows AFD to correctly establish a new set point following a temperature experience in order to continue to respond bidirectionally with high sensitivity to temperature changes. A feature of many sensory systems is the rapid evolution of sensory receptors that enable animals to adapt to their specialized environmental niche (Bear et al., 2016; Baldwin and Ko, 2020; Yau and Hardy, 2009; Julius and Nathans, 2012). The functioning of sensory systems is further optimized by the presence of species- and modality-specific molecules and mechanisms (Zang and Neuhauss, 2018; Huber, 2001). The transient upregulation of *pyt-1* during a specific phase of AFD response plasticity may be critical for fine-tuning AFD responses possibly via modulation of one or more signaling proteins required for production and/or hydrolysis of cGMP. We propose that species-specific evolution and neuron-specific expression of molecules such as PYT-1, along with acquisition of stimulus-dependent gene regulatory architecture, may provide an elegant mechanism by which sensory transduction mechanisms can be flexibly yet specifically modified.

The roles of regulatory molecules including kinases such as CaMKIV, transcription factors such as CREB and MEF2, as well as subsets of downstream target genes such as Bdnf and Arc in driving experience-dependent neuronal plasticity have largely been studied using gene knockout models (Bourtchuladze et al., 1994; Ernfors et al., 1994; Jones et al., 1994; Shepherd et al., 2006). A confounding factor in these analyses has been the inability to distinguish between possible roles of these molecules in regulating basal vs activity-regulated properties of the expressing neurons. Activity responses can be teased apart from basal expression by identifying and disrupting endogenous activity-dependent cis-regulatory regions, allowing for specific testing of their functional consequences (Hong et al., 2008). The similar response defects of *pyt-1* single CRE mutants and *pyt-1* protein null mutants at 4 hrs after a temperature upshift confirm the critical importance of temperature and CREB-mediated upregulation of this gene in mediating a defined aspect of AFD response plasticity but not other AFD response properties. The ability to specifically target the cis-regulatory site(s) driving activity-dependent gene expression in vivo provides a powerful method to dissect the contributions of individual genes to defined aspects of neuronal and behavioral plasticity. While the experience-dependent gene expression changes described here are controlled by signaling pathways and transcriptional mechanisms that are common to many neurons, these genes appear to be temperature-regulated only in AFD and play specific roles in regulating AFD properties. Processes that specify a neuron’s static molecular identity may be permissive for their ability to modify that identity in response to encountered stimuli (Hobert and Kratsios, 2019). Thus, specialized activity-dependent gene batteries may be a feature of subsets of neuron types that can only be detected via the use of defined stimuli and characterization of single cell transcriptomes (Yap et al., 2021; Mardinly et al., 2016; Tsukahara et al., 2021; Gray and Spiegel, 2019). A major future challenge will be to comprehensively identify and characterize stimulus-specific gene expression programs at single neuron resolution in vivo, and establish how changes in the expression of single genes and gene batteries encode the animal’s experience to shape defined aspects of its physiological and behavioral responses.

## ACKNOWLEDGEMENTS

We are grateful to Xicotencatl Gracida for help with the TRAP protocol, and Nandini Mani and Radhika Subramanian for assistance with sequence analysis of PYT-1. We thank the Caenorhabditis Genetics Center and the National BioResource Project for strains, and Daniel Colón-Ramos for reagents. We thank members of the Sengupta lab, Bob Datta, Oliver Hobert, Steven Flavell, and Jenna Sternberg for advice and critical comments on the manuscript. This work was funded in part by the NIH (R35 GM404386 – P.S., F32 NS112453 and T32 NS007292 – N.H. T32 GM139798 – S.B., T32 MH019929 – J.S.), Ministry of Science and Technology of China (2021ZD0202603 – Y.V.Y.) and the Natural Sciences and Engineering Research Council of Canada (NSERC) and the Canadian Institutes of Health Research (CIHR) (J.A.C.)

## Methods

### C. elegans strains and genetics

Worms were grown at 20°C on nematode growth media (NGM) plates seeded with E. coli OP50 unless noted otherwise. The wild-type strain used was C. elegans variety Bristol strain N2. Transgenic animals were generated using experimental plasmids at 3-100 ng/ µl and the coinjection marker at 30-50 ng/ µl. Strains containing two or more alleles were generated using standard genetic crosses. The presence of specific molecular lesions and genomic edits was confirmed by sequencing. To stably integrate *ttx-1p::tagRfp* and *ttx-1p::egfp::rpl-1a* sequences into the genome, young transgenic adults carrying extrachromosomal arrays were irradiated with 300 µJ UV light. Animals homozygous for the integrated array were backcrossed six times prior to use. A complete list of all strains used in this work is provided in Supplementary file 1.

### Plasmid construction

Promoter sequences, genomic loci, and cDNAs were amplified from genomic DNA or plasmids, or a C. elegans cDNA library generated from a population of mixed stage animals, respectively (Nechipurenko et al., 2016). Plasmids were constructed using standard restriction enzyme cloning or Gibson assembly (New England BioLabs). All plasmids used in this work are described in Supplementary file 2.

### CRISPR/Cas9-based genome engineering

All crRNAs, tracrRNAs and Cas9 protein were obtained from Integrated DNA technologies (IDT). All gfp insertions were made immediately before the stop codon of each relevant gene in the N2 genetic background.

*Reporter-tagged alleles: gcy-18(oy165[gcy-18::gfp]*: A donor plasmid was created by first cloning 2 kb genomic DNA surrounding the insertion site into pMC10 (gift of M. Colosimo), followed by addition of AarI restriction sites via site directed mutagenesis (Agilent) to enable insertion of gfp sequences. The gfp sequence was inserted using a crRNA (5’-ctgaatgtagtttgtagtcg-3’) and a gfp donor with 1 kb homology arms amplified from the donor plasmid. *ins-39(oy167[ins-39::SL2::gfp])*: a crRNA (5’-gatggagcattgatcagagc-3’) and a donor containing SL2::gfp amplified using oligonucleotides containing 35 bp homology arms were used to insert SL2::gfp sequences (Servello et al., 2022). *dac-1(oy172[dac-1::gfp])*: A crRNA (5’-gtcttcggacggaaattctg-3’) and a donor containing gfp sequences amplified using oligonucleotides containing 35 bp homology arms were used to insert gfp sequences. F08H9.4(syb5551[F08H9.4::gfp]) was obtained from SunyBiotech. pyt-1(oy169[pyt-1::gfp]): A crRNA (5’-tacatgacgtcatcatatga-3’) and a gfp donor with 400 bp homology arms amplified from a GFP-encoding plasmid were used to insert gfp sequences.

*Deletion allele:* The *pyt-1(oy160)* deletion allele was generated according to published protocols (Ghanta et al., 2021). 2 crRNAs (5’-tagaaaagtgaattcactgt-3’ and 5’-tggaatgaaacgttgactgc-3’) targeting sequences upstream and downstream of the *pyt-1* coding region and an ssODN donor (5’-cttctaatacgcataaagtcagtatttatagttcagaagtaaatatatttgatttgtgtttag ttttttc-3’) containing 35 bp of homology 5’ and 3’ to the cut sites were injected along with Cas9 protein. The injection mix contained: ssODN donor (110 ng/ µl), crRNAs (28 ng/ µl each), tracrRNA (100 ng/ µl), Cas9 (250 ng/ µl), and co-injection marker (myo-3p::mCherry (50 ng/ µl)). The progeny of transgenic animals was subsequently examined for the presence of the desired deletion via sequencing.

*Mutations in predicted CREs:* To mutate a predicted CRE upstream of *pyt-1(oy169)*, a crRNA (5’-tttgtgacgtcgtctgcaaa-3’) and an ssODN donor (5’-ttttcttctcagtacttgagccaataaaccttttgaaggtttgtctgcaaatccataccaaatctgccaccacaaca-3’) were used to generate *pyt-1(oy169 oy173)*. To mutate CREs upstream of *dac-1(oy172)*, two crRNAs (5’-gatatttttcccagaaagct-3’ and 5’-tctattagatgacgctcatg-3’) and an amplified donor containing mutations in two predicted CREs (tgacgtag>aaggtttg and gtacgtca>aaggtttg) were used to replace the sequences between the crRNA-targeted cut sites and generate *dac-1(oy172 oy175)*. In the process of generating *dac-(oy172 oy175)*, animals carrying a deletion between these two cut sites was also isolated, yielding the allele *dac-1(oy172 oy174)*.

### Translating Ribosome Affinity Purification (TRAP)

TRAP was performed according to published protocols (Gracida and Calarco, 2017) with the following modifications. Worms were cultured on 10X concentrated E. coli HB101 seeded at 2 ml per 15 cm plate. Each plate contained 50,000 growth-synchronized 1-day old adult animals. Animals were growth synchronized by bleaching adults, collecting eggs, and then arresting hatched L1s in the absence of food for 16 hours. A total of 10 plates was used per sample. Worms were cultivated at 20°C until the L4 stage when animals from the same population were placed on plates at 15°C or 25°C overnight. Worm collections were completed within 15 mins (from incubator to flash freezing) to minimize temperature variations. Each immunoprecipitation used worm lysate containing 2000-3000 µg of total RNA per sample. As validation for enrichment of AFD RNA by immunoprecipitation, gcy-8 mRNA was detected using a OneStep RT-PCR kit (QIAGEN) from 3 ng of template RNA. All TRAP experimental steps were performed on samples grown at the two temperatures in parallel using the same TRAP affinity matrix.

### RNA-Sequencing

10 ng RNA from each worm sample was used as input for cDNA synthesis using a SMART-Seq v4 Ultra Low Input RNA kit (Takara). Sequencing libraries were prepared from 500 pg of cDNA with a Nextera XT DNA Library Prep kit (Illumina). Libraries were paired-end sequenced at 75×75 bases on a NextSeq 500 system (Illumina). Sequencing reads were adapter trimmed using cutadapt (Martin, 2011) with the following options: quality cutoff 20, –trim-n, –minimum-length=50, and then mapped to the C. elegans genome (WBcel235/ce11), and counted using STAR (Dobin et al., 2013) with –quantMode GeneCounts. Differential expression analysis was performed in R with DESEQ2 (Love et al,. 2014) (GEO accession number GSE222226). The volcano plot was generated in R using EnhancedVolcano (https://github.com/kevinblighe/EnhancedVolcano). Gene set enrichment analyses were performed in R using the clusterProfiler package (Yu et al., 2012).

### Thermotaxis behavior

Thermotaxis behaviors were performed as previously described (Luo et al., 2014) with the following modifications. Well-fed animals were grown at 20°C to the L4 stage and then cultured at 15°C overnight prior to performing negative thermotaxis assays. For positive thermotaxis assays, animals cultured at 15°C were shifted to 25°C for four hours prior to the assay. The temperature gradient in the assay arena was either 18-22°C (for negative thermotaxis) or 19-23°C (for positive thermotaxis) at a steepness of 0.18°C/cm (Ji et al., 2021). Temperature was controlled by a Peltier system [colder side; H-bridge amplifier (Accuthermo FTX700D), PID controller (Accuthermo FTC100D), Peltier (DigiKey)] and heater system [warmer side; PID controller (Omega CNi3244), solid-state relay Omega SSRL240DC25), and cartridge heaters (McMaster-Carr 3618K403)]. A 22.5 cm square NGM agar pad was used for the assay, and the temperature of agar edges and center were confirmed with a digital thermometer (Fluke Electronics) prior to each assay. 15-25 worms in M9 buffer were placed at the center of the gradient at the start of the assay. Animal movement was imaged at 2 fps for 60 min using a Mightex camera (BTE-5050-U). Animal trajectories were detected and analyzed using custom LabView (National Instruments) and MATLAB (Mathworks) scripts (Gershow et al., 2012) (https://github.com/samuellab/MAGATAnalyzer). All assays were performed using one day-old adults.

### Calcium imaging

Temperature-evoked calcium dynamics in AFD were measured essentially as described previously (Takeishi et al., 2020; Takeishi et al., 2016; Yu et al., 2014) with the following modifications. Animals were cultivated at 20°C until the L4 stage and then shifted to the indicated temperatures. One day-old well-fed adults were immobilized in 10 mM tetramisole on an agarose pad (5% in M9 buffer) on a cover glass, and mounted under a second cover glass for imaging. The sample was transferred to a Peltier temperature control system on the microscope stage. Animals were subjected to linear temperature ramps rising at 0.05°C/s via temperature-regulated feedback using a temperature controller (Accuthermo FTC200), an H-bridge amplifier (Accuthermo FTX700D), and a thermistor (McShane TR91-170). Videos of calcium dynamics at the AFD sensory endings were captured using a Zeiss 40X air objective (NA 0.9) or a Zeiss 10X air objective (NA 0.3) on a Zeiss Axioskop2 Plus microscope, using a Hamamatsu Orca digital camera (Hamamatsu), and MetaMorph software (Molecular Devices). Data were analyzed using custom scripts in MATLAB (Mathworks) (https://github.com/wyartlab/Cantaut-Belarif-et-al.-2020) (Sternberg et al,. 2018; Cantaut-Belarif, 2020). T*AFD was calculated as described previously (Takeishi et al., 2016).

### cGMP imaging

An AFD-expressed FlincG3 cGMP sensor (Woldemariam et al., 2019) was used to measure changes in cGMP concentration. cGMP imaging was performed essentially as described above for calcium imaging. FlincG3 fluorescence was observed to decrease substantially during imaging due to photobleaching. For visualization, the deltaF/F traces were fit with an exponential curve which was subtracted to generate the data in Figure 7F.

### Analyses of GFP fluorescence

To visualize and quantify GFP fluorescence in animals expressing reporter-tagged alleles, well-fed one day-old adult animals grown under indicated temperature conditions were immobilized with 20 mM tetramisole, mounted on 10% agarose pads on slides, and imaged either on a Zeiss Axio Imager M2 epifluorescent microscope with a 63x oil objective (NA 1.4) or on a Zeiss Axio Observer with a Yokogawa CSU-X1 spinning disk confocal head (3i Marianas system) with a 63x oil objective (NA 1.4). AFD was typically identified by co-expression of the AFD-specific marker ttx1p::tagRfp (Satterlee et al., 2001). Images for quantification were acquired with no red marker in the background. Images were processed in ImageJ, and expression was quantified from a maximum projected z-stack as corrected total cell fluorescence (CTCF) using the equation CTCF = Integrated Density – (Area of selected cell ROI x Mean fluorescence of a nearby background ROI). Protein localization/colocalization at AFD sensory endings was imaged using a Zeiss LSM880 AiryScan Fast Confocal System in the AiryScan configuration with a 63x oil objective (NA 1.4). Animals expressing daf-7p::gfp (ksIs2) and daf-28p::gfp (mgIs40) were cultivated at 27°C under the same conditions used for performing dauer assays (see below) and imaged as L1s 20-24 hrs after egg laying. Imaging conditions were identical to those used for AFD-expressed reporters. ASI and ASJ neurons were identified by soma position.

### Single molecule fluorescent in situ hybridization (sm-FISH)

The animals used were *pyt-1(oy169[pyt-1::gfp]); Ex[ttx-1p::tagRfp]*. Growth synchronized L1 larvae were cultivated at 20°C until the L3 stage. Worms were then shifted to different temperatures as indicated and fixed the next day as L4s. L4s were used because the thicker cuticle of 1 day old adults decreased staining efficacy. Stellaris gfp RNA FISH probes conjugated to Quasar 670 (Biosearch Technologies) were used. The Stellaris smFISH protocol was followed with the following modifications. After fixation, worms were permeabilized in 70% ethanol for 2 nights at 4°C. Hybridization was performed with probe concentration of 250 nM at 37°C overnight. No nuclear counterstain was used. Before mounting onto slides, samples were washed once with 2x SSC, then once with GLOX buffer, then suspended in GLOX buffer containing glucose oxidase and catalase. The sample was immediately used for imaging. Images were acquired on a Zeiss Axio Imager M2 epifluorescent microscope with a 63x oil objective (NA 1.4). z-stacks were acquired at 0.5 µm thickness. z-stacks were analyzed in ImageJ. AFD was identified using the AFD-specific *ttx-1p::tagRfp transgene*. No Quasar 670 signal was observed outside of AFD. Background subtraction was performed on the Quasar 670 channel, and puncta were quantified by scanning through the z-slices.

### Dauer formation assays

Dauer formation assays were performed as described previously (Neal et al., 2013; O’Donnell et al., 2018), with the following modifications. Assay plates were seeded with 10 µl of 16 mg/ml OP50. Live or heat-killed OP50 was used for 27°C-induced or pheromone-induced assays, respectively. Growth-synchronized adult hermaphrodites were allowed to lay 50-75 eggs per assay plate and plates were examined for dauers 70 hrs later. Pheromone-induced dauer formation assays were performed at 25°C using the ascr5 pheromone (Kaplan et al., 2011) at a final concentration of 600 nM in the plate agar. Dauer larvae were identified by their characteristic morphology.

### Motif enrichment analysis and identification of putative CREs

Transcription factor binding motifs represented as position weight matrices (PWMs) were acquired from a collection of CIS-BP database PWMs curated by MEME (Weirauch et al., 2014). The PWM set used in this analysis contained all available PWMs obtained for human, mouse, worm, and fly transcription factors (total of 2173 PWMs). This PWM set was then used as input for the SEA enrichment algorithm (Bailey and Grant, 2021) provided by the MEME suite of sequence analysis tools (Bailey et al., 2015), together with 500 bp DNA sequences immediately upstream of the start codons of each AFD-specific temperature-upregulated gene. Promoter sequences upstream of AFD-specific temperature-insensitive genes were used as control sequences. Enrichment by SEA was run with default parameters. SEA returned a list of 57 motifs that were over-represented in temperature-upregulated versus temperature-insensitive promoters. These motifs were then clustered by similarity into families of similar motifs using the matrix-clustering algorithm from RSAT (Castro-Mondragon, 2017) with the following parameters: –cor_th 0.7 –Ncor_th 0.5. To identify individual CREs, the promoters of a subset of temperature-upregulated genes were scanned using FIMO (Grant et al., 2011) for matches to any of the enriched CRE motifs identified by SEA. Highest scoring matches were chosen for perturbation via gene editing.

### Quantification and analyses

Summary plots of RNA-Seq data were generated with their associated packages in R, except for the plot of TRAP-Seq AFD expression versus CeNGEN AFD gene expression (Figure 1B), which was plotted in Prism 6 (GraphPad). Plots of fluorescence intensity, smFISH RNA puncta, dauer quantifications, T*AFD, and thermotaxis behavior quantifications were generated with Prism 6. GCaMP/FlincG3 fluorescence traces and thermotaxis assay example images were generated with MATLAB (Mathworks). Example images of fluorescent expression/localization reporters were made with ImageJ. The phylogenetic tree (Figure S5A) was generated with Geneious Prime (Biomatters). Multiple sequence alignment plots (Figure S3C, Figure S5B) were generated with Jalview 2.11.2.5 (Waterhouse et al., 2009). Statistical analyses were performed in Prism 6. Statistical test details, the number of analyzed samples and biologically independent experiments are reported in each figure legend.

## Code availability

All code used in this study is available at https://github.com/SenguptaLab/AFD_gene_expression.

---

**Figure S1.**
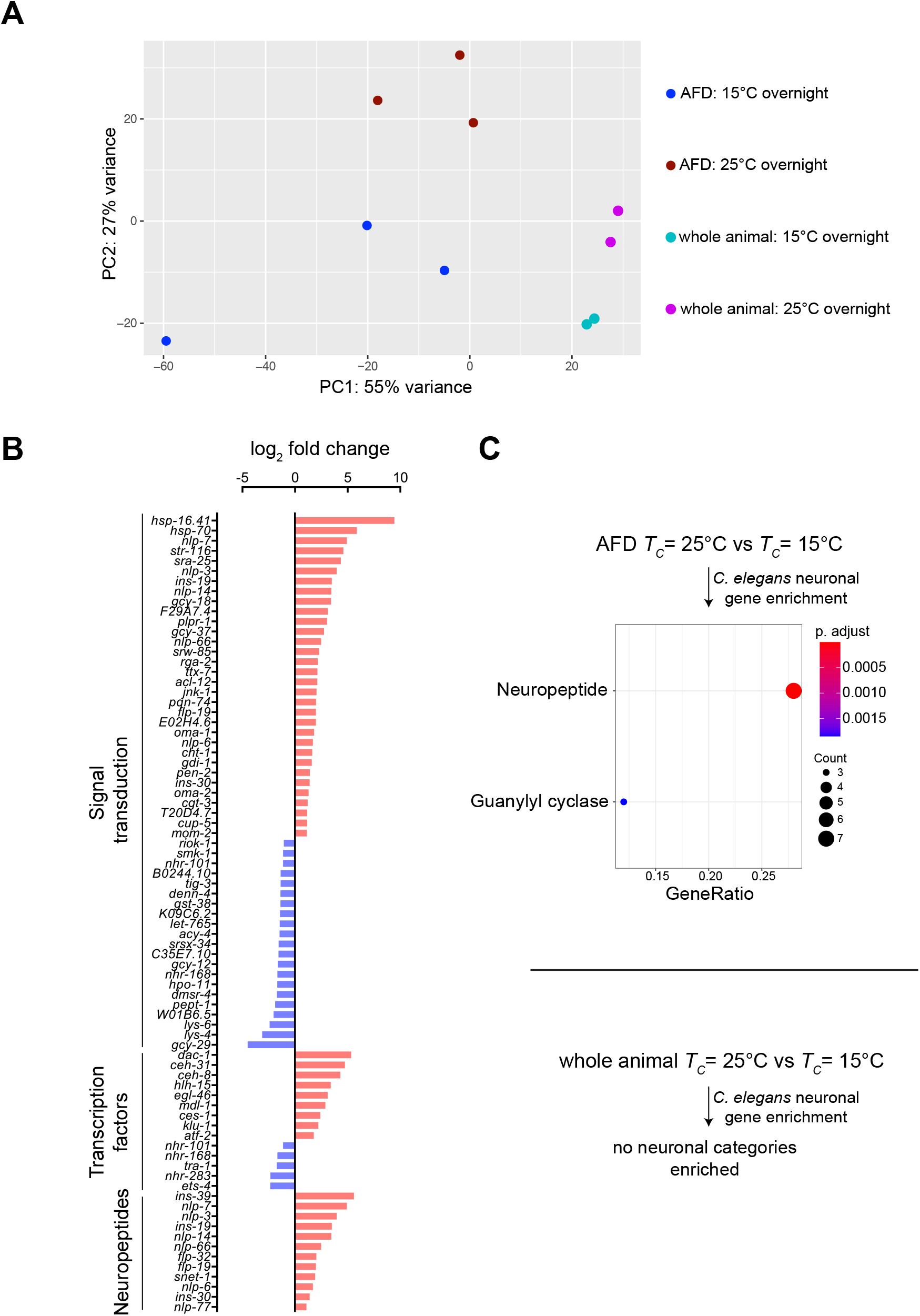
Analysis of TRAP-Seq data (Related to Figure 1). **A)** DESeq2-generated principal component analysis plot of TRAP-Seq data clustered by tissue and temperature cultivation condition. Each dot represents a single replicate. **B)** Waterfall plot of temperature-dependent expression of genes in AFD encoding selected protein classes. Displayed genes exhibit > 2 or < -2 log2 fold change with a threshold of adjusted p-value < 0.05. Genes in the “Signal transduction” category were filtered by the gene ontology term “signal transduction” – GO:0007165. The categories “Transcription factors” and “Neuropeptides” were derived from a curated list of *C. elegans* transcription factors and neuronal genes (Hobert, 2013; Valperga and de Bono, 2022). **C)** Enriched neuronal gene classes from temperature-dependent genes in AFD and whole animal samples using clusterProfiler (Yu et al., 2012) on curated *C. elegans* neuronal genes (Hobert, 2013; Valperga and de Bono, 2022).

**Figure S2.**
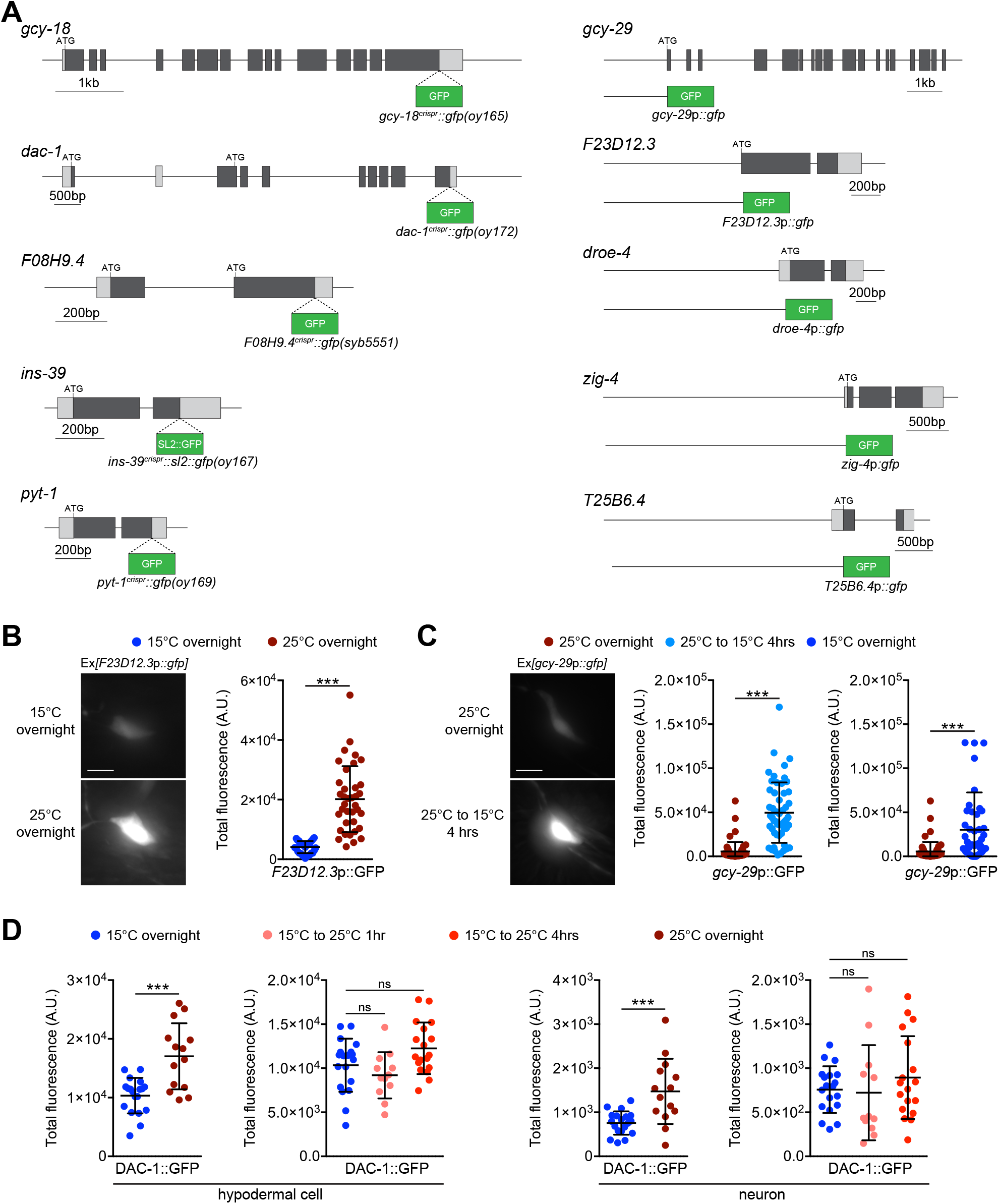
Temperature-regulated expression of reporter fusion genes in AFD (Related to Figure 2). **A)** (Left) Genomic structures of endogenously reporter tagged genes examined in Figure 2. (Right) Structures of promoter::GFP fusion reporters for an additional subset of genes. Expression from these genes were examined from extrachromosomal arrays in transgenic animals (Table S1). Light and dark gray boxes indicate untranslated regions and coding DNA, respectively. **B**,**C)** (Left) Representative images of *F23D12*.3p::GFP (B) and *gcy-29*p::GFP (C) expression in AFD soma of adult animals grown at the indicated conditions. Scale bar: 5 µm. (Right) Quantification of GFP levels in adult animals grown at the indicated conditions. Each dot is a measurement from a single AFD neuron. Horizontal and vertical lines indicate mean and SD, respectively. n = 33-55 neurons from at least two biologically independent experiments. **D)** Quantification of DAC-1::GFP levels in a hypodermal cell (left) or unidentified head neuron in adult animals grown at the indicated conditions. Each dot is a measurement from a single cell. Horizontal and vertical lines indicate mean and SD, respectively. n = 12-19 cells from at least two biologically independent experiments. For all panels, *** indicates different at p<0.001 (t test or one-way ANOVA with Dunnett’s multiple comparisons correction); ns – not significant.

**Figure S3.**
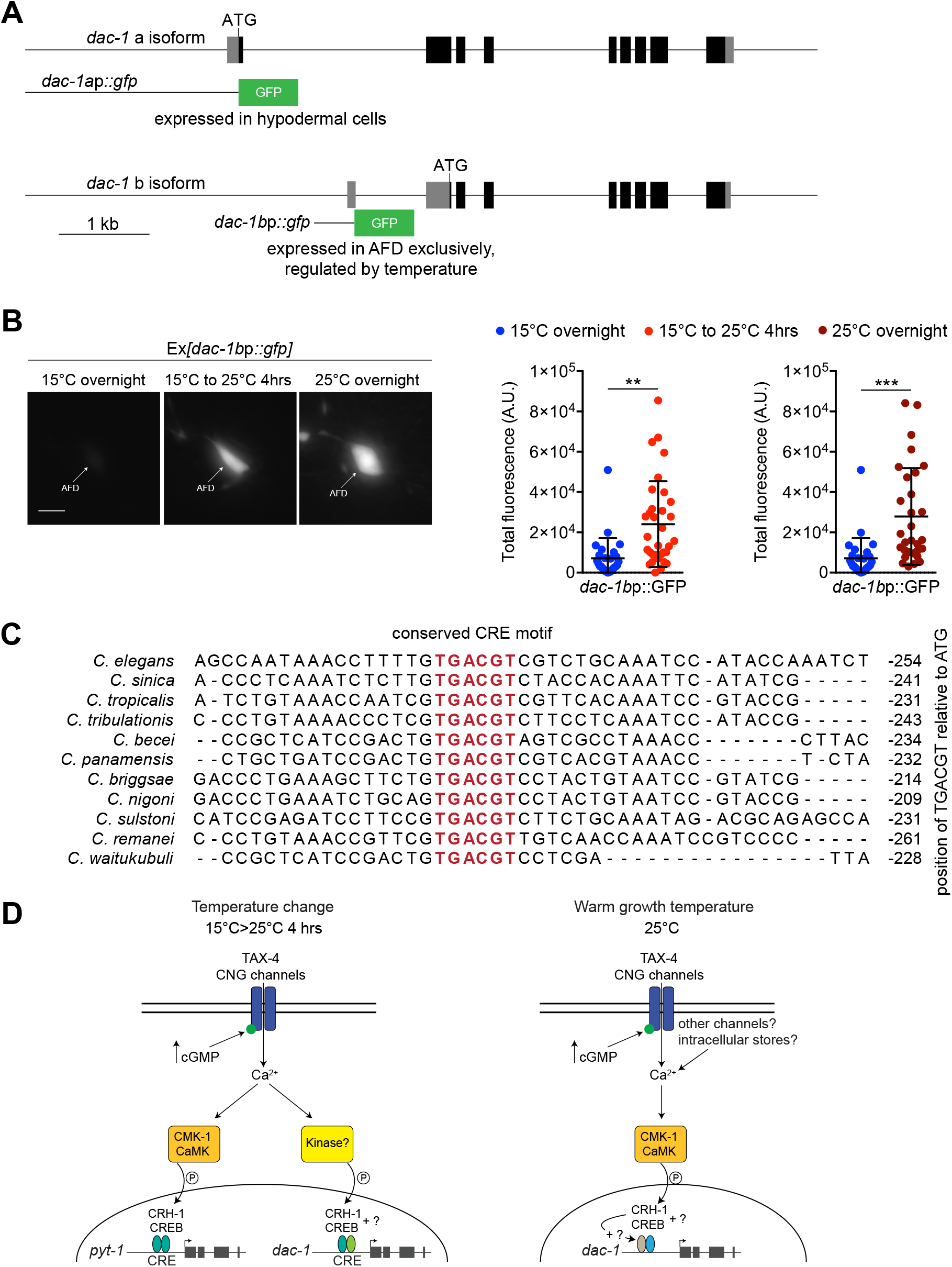
Analysis of *cis*-regulatory sequences upstream of *pyt-1* and *dac-1* (Related to Figure 5). **A)** *dac-1* genomic locus showing the a and b isoforms. Grey and black boxes indicate untranslated regions and coding DNA, respectively. The structures of the transcriptional reporter genes examined are shown below. **B)** (Left) Representative images of GFP driven by the *dac-1b* promoter in adult animals grown at the indicated conditions. Scale bar: 5 µm. (Right) Quantification of *dac-1b*p::GFP levels in adult animals grown at the indicated conditions. Each dot is a measurement from a single AFD neuron. Horizontal and vertical lines indicate mean and SD, respectively. n = 27-33 neurons from at least two biologically independent experiments. All conditions were assayed in parallel; data from 15°C overnight grown animals are repeated in the two panels. ** and *** indicate different at p<0.01 and 0.001, respectively (one-way ANOVA with Dunnett’s multiple comparisons correction). **C)** Conservation of a predicted CRE motif (red) in the upstream regulatory sequences of a subset of *pyt-1* orthologs in other nematodes. Sequences were acquired from WormBase ParaSite BLAST (https://parasite.wormbase.org/). Sequences were aligned with Clustal Omega (https://www.ebi.ac.uk/Tools/msa/clustalo/). Formatting was performed with Jalview. **D)** Summary of signaling pathways translating temperature stimuli into changes in expression of *dac-1* and *pyt-1*.

**Figure S4.**
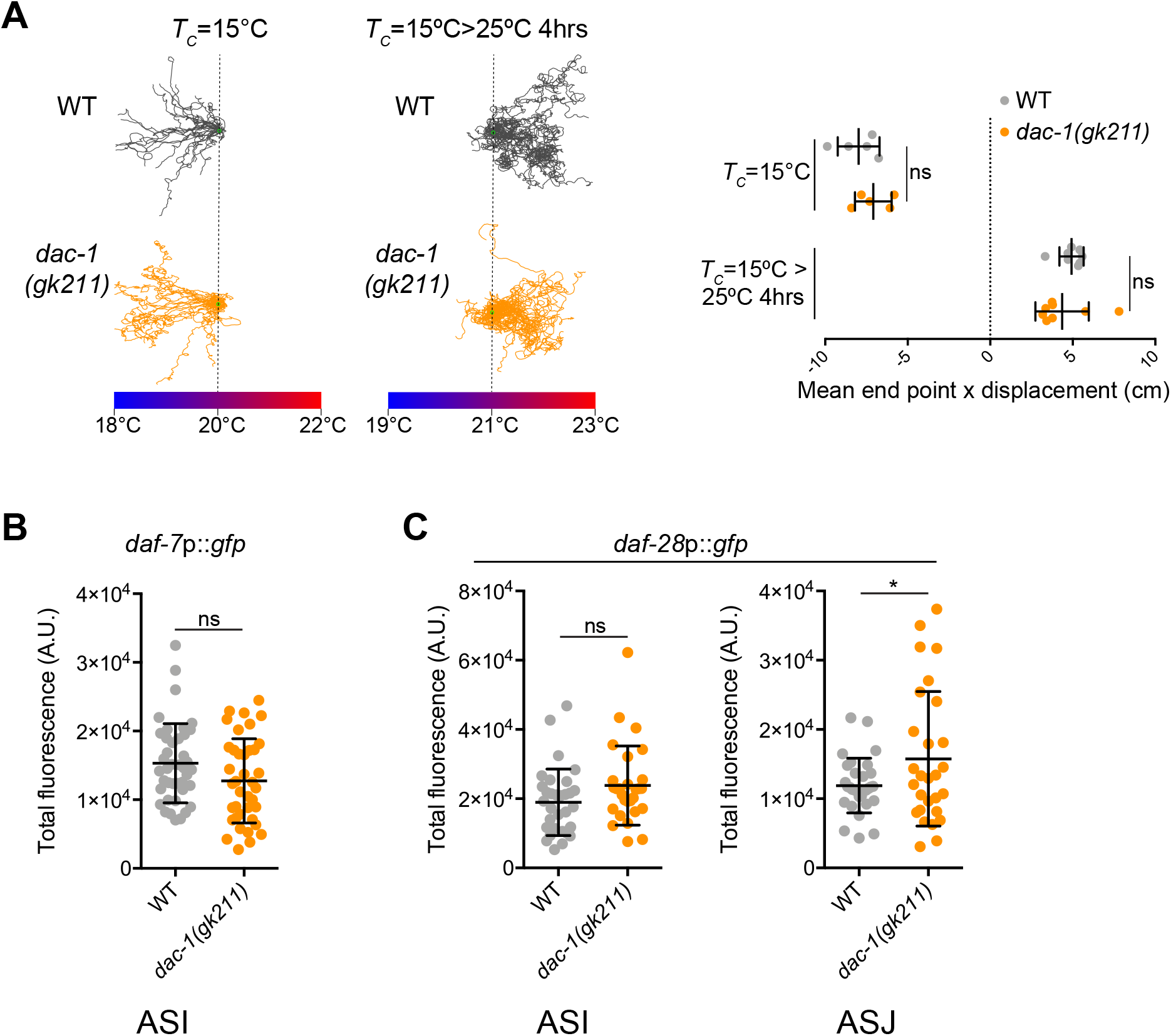
DAC-1 does not regulate DAF-7 TGF-β or DAF-28 insulin in ASI or ASJ (Related to Figure 6). **A)** (Left) Representative thermotaxis behavior trajectories of adult wild-type and *dac-1(gk211)* animals in a single assay on thermal gradients grown at the indicated conditions. Vertical dashed lines indicate starting temperature on the gradient. (Right) Quantification of thermotaxis assay end points relative to starting position. Each dot indicates the average end point distance of all animals in a single assay. Vertical and horizontal lines indicate mean and SD, respectively. n = 5-8 assays of 15-25 animals each from at least two independent days. **B**,**C)** Quantification of *daf-7*p::GFP (B) and *daf-28*p::GFP (C) in ASI and/or ASJ neurons of L1 larvae grown at 27°C. Each dot is a measurement from a single neuron. Horizontal and vertical lines indicate mean and SD, respectively. n = 28-42 neurons from at least two biologically independent experiments. * indicates different at p<0.05 (t test); ns – not significant.

**Figure S5.**
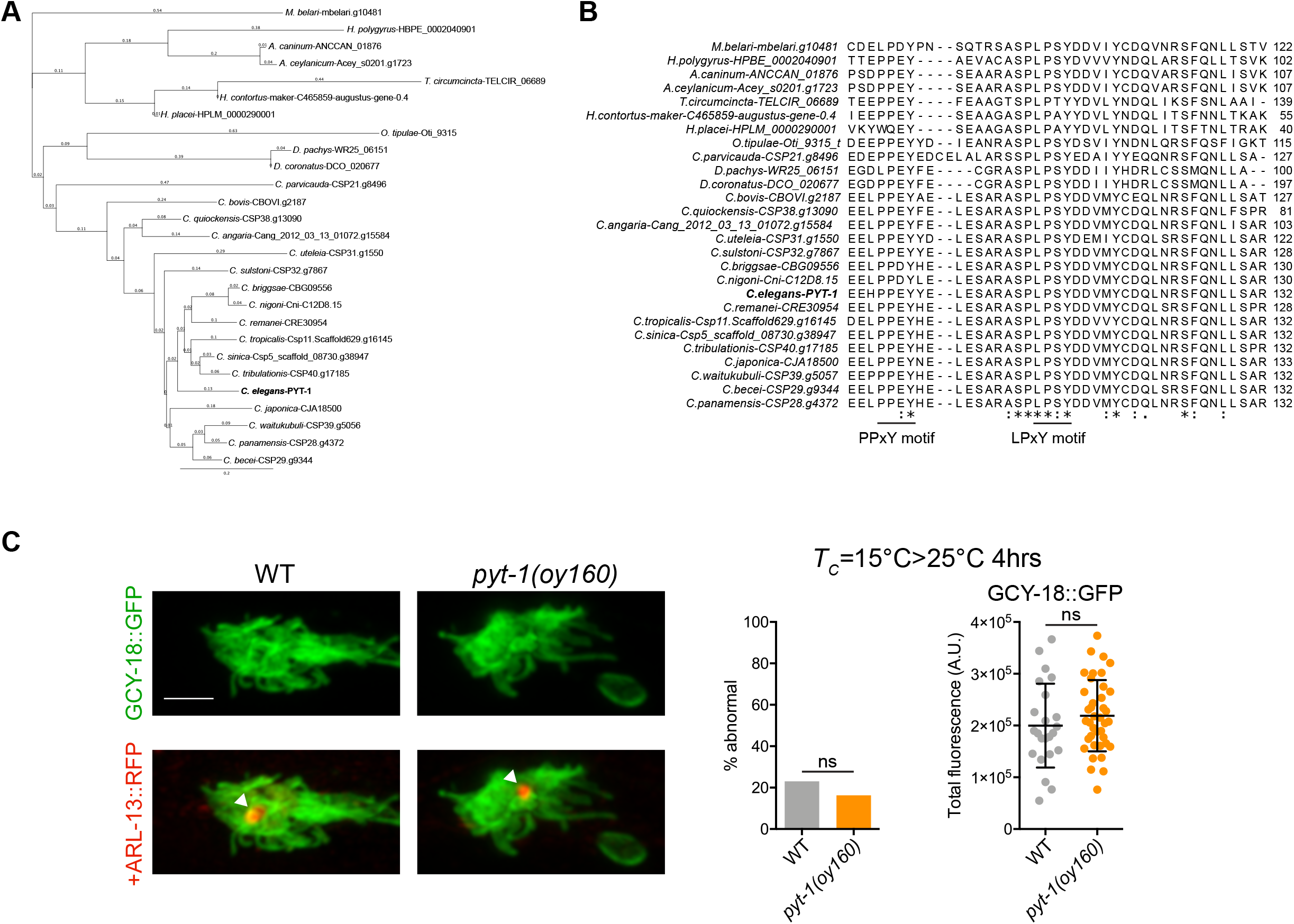
PYT-1 is conserved in a subset of nematodes and contains conserved PY motifs (Related to Figure 7). **A)** Phylogenetic tree of PYT-1 and orthologs. PYT-1 orthologs and their sequences were acquired from WormBase ParaSite BLAST (https://parasite.wormbase.org/). Sequences were aligned and the phylogenetic tree was constructed with Clustal Omega (https://www.ebi.ac.uk/Tools/msa/clustalo/). Formatting was performed with Geneious Prime. **B)** Conservation of the PPxY and LPxY motifs in the C-terminal domains of PYT-1 and orthologs in other nematodes. **C)** (Left) Representative images of endogenously tagged GCY-18::GFP and *gcy-8*p::ARL-13::RFP localization at the AFD sensory endings. The cilium is indicated with an arrowhead. Scale bar: 2 µm. Anterior is at left. (Middle) Quantification of AFD sensory ending morphology in adult animals from the indicated genotypes grown at 15°C overnight and then shifted to 25°C for 4 hours before imaging. AFD sensory endings were visualized with endogenously tagged GCY-18::GFP. n = 39-49 animals. (Right) Quantification of GCY-18::GFP levels in adult animals from the indicated genotypes grown at 15°C overnight and then shifted to 25°C for 4 hours before imaging. Each dot is a measurement from a single AFD sensory ending. Horizontal and vertical lines indicate mean and SD, respectively. n = 23-39 sensory endings from at least two biologically independent experiments. ns – not significant (Fisher’s exact test or t test).

**Figure S6.**
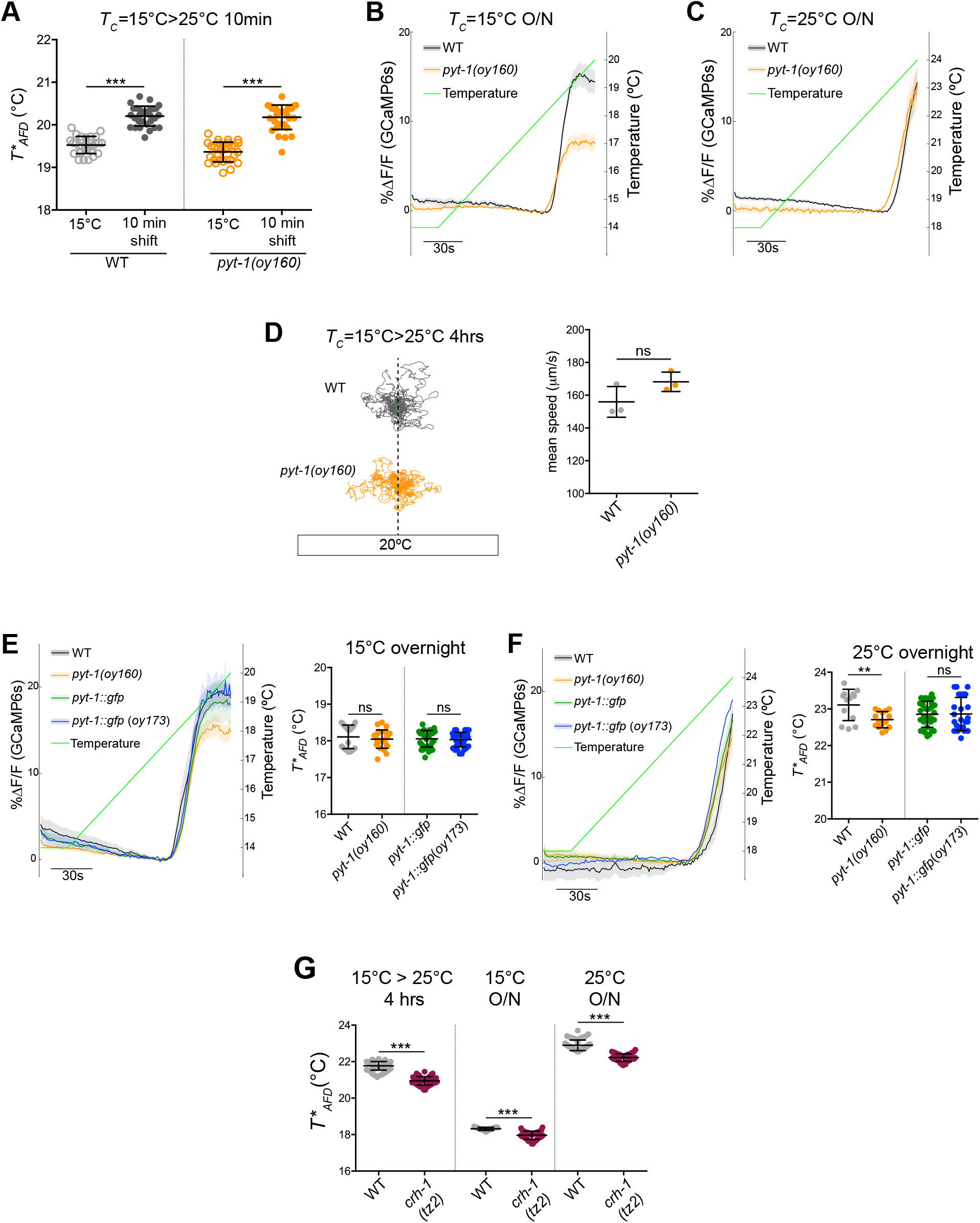
CREB-mediated upregulation of PYT-1 is necessary only under specific temporal conditions to regulate neuronal plasticity (Related to Figure 7) **A)** Quantification of *T*AFD* measured via GCaMP imaging at the AFD sensory endings in wild-type and *pyt-1* animals at the indicated temperature conditions. Each dot is a measurement from a single AFD neuron. Horizontal and vertical bars are the mean and SD, respectively. n = 29-31 animals from at least two biologically independent experiments. **B**,**C)** GCaMP traces from AFD in adult animals during the temperature ramp protocol (green line). Thick lines and shading indicate the average ΔF/F change and SEM, respectively. These traces come from the 15°C or 25°C O/N condition quantified in Figure 7E. **D)** Representative trajectories of wild-type and *pyt-1* mutants in a single assay on an isothermal plate at 20°C. Animals were grown under temperature conditions used for positive thermotaxis (see Figure 7G). n = 3 assays of 15-25 animals. **E**,**F)** (Left) GCaMP traces from AFD in adult animals grown at the indicated conditions during the temperature ramp protocol (green line). Thick lines and shading indicate the average ΔF/F change and SEM, respectively. (Right) Quantification of *T*AFD* for the indicated genotypes grown at the indicated conditions. Each dot is a measurement from a single animal. Horizontal and vertical lines indicate mean and SD, respectively. n = 13-37 animals from at least two biologically independent experiments. **G)** Quantification of *T*AFD* from animals of the indicated genotypes grown at the indicated conditions. Each dot is a measurement from a single animal. Horizontal and vertical lines indicate mean and SD, respectively. n = 17-55 animals from at least two biologically independent experiments. For all panels, ** and *** indicate different at p<0.01 and 0.001, respectively (t test,); ns – not significant.

**Table S1.**
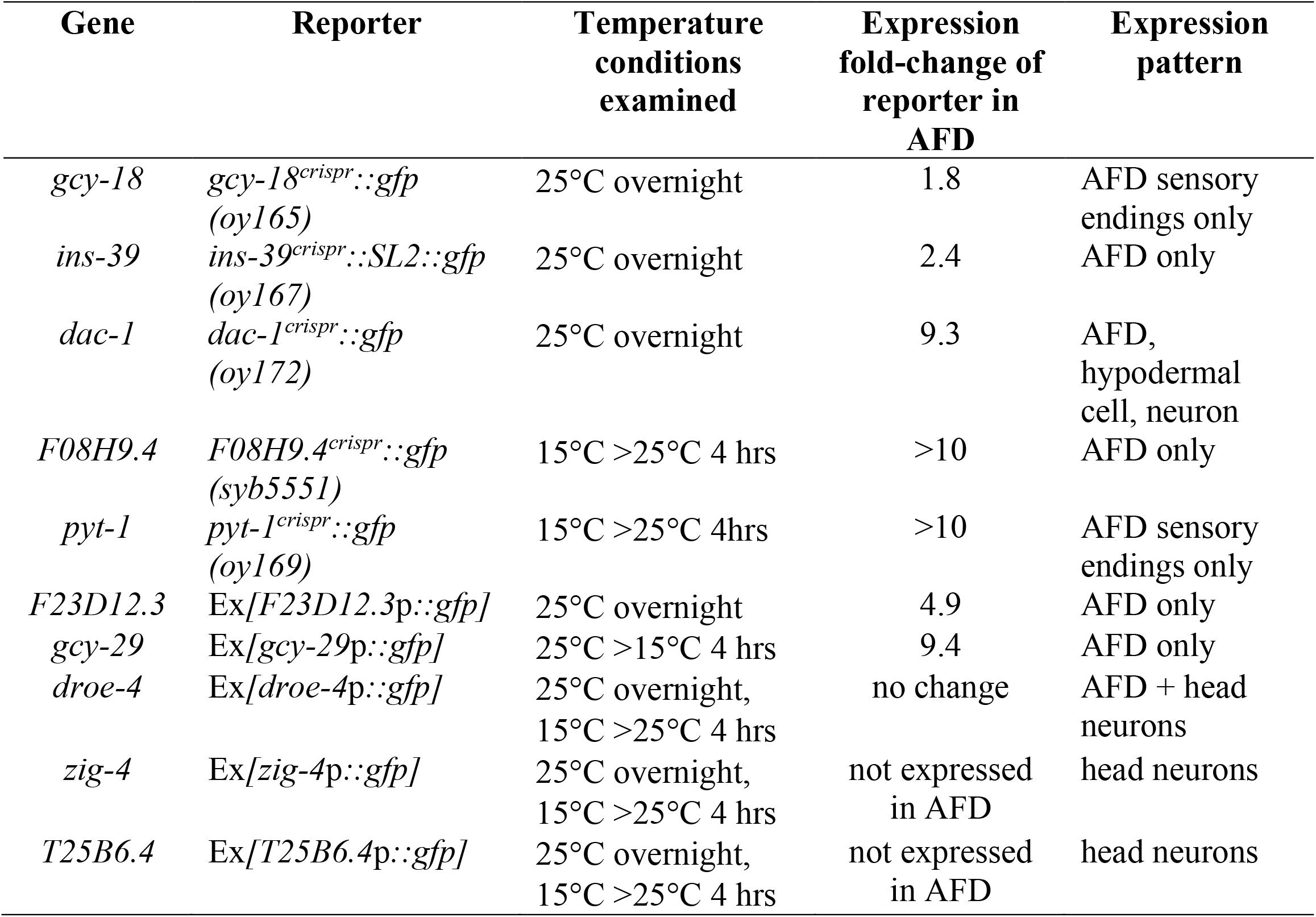
related to Figure 2. Temperature-regulated expression changes of examined reporter genes.

**Table S2.**
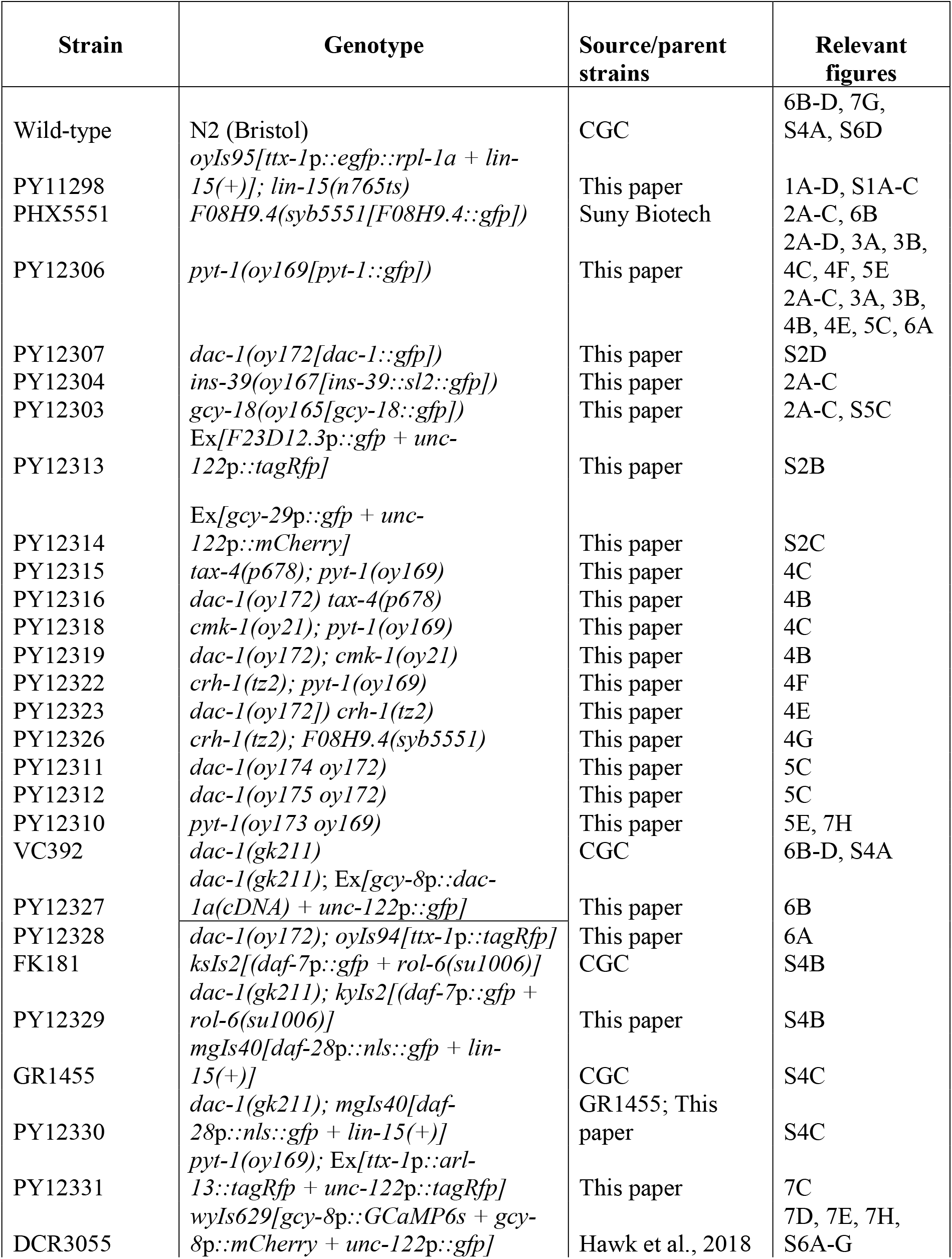
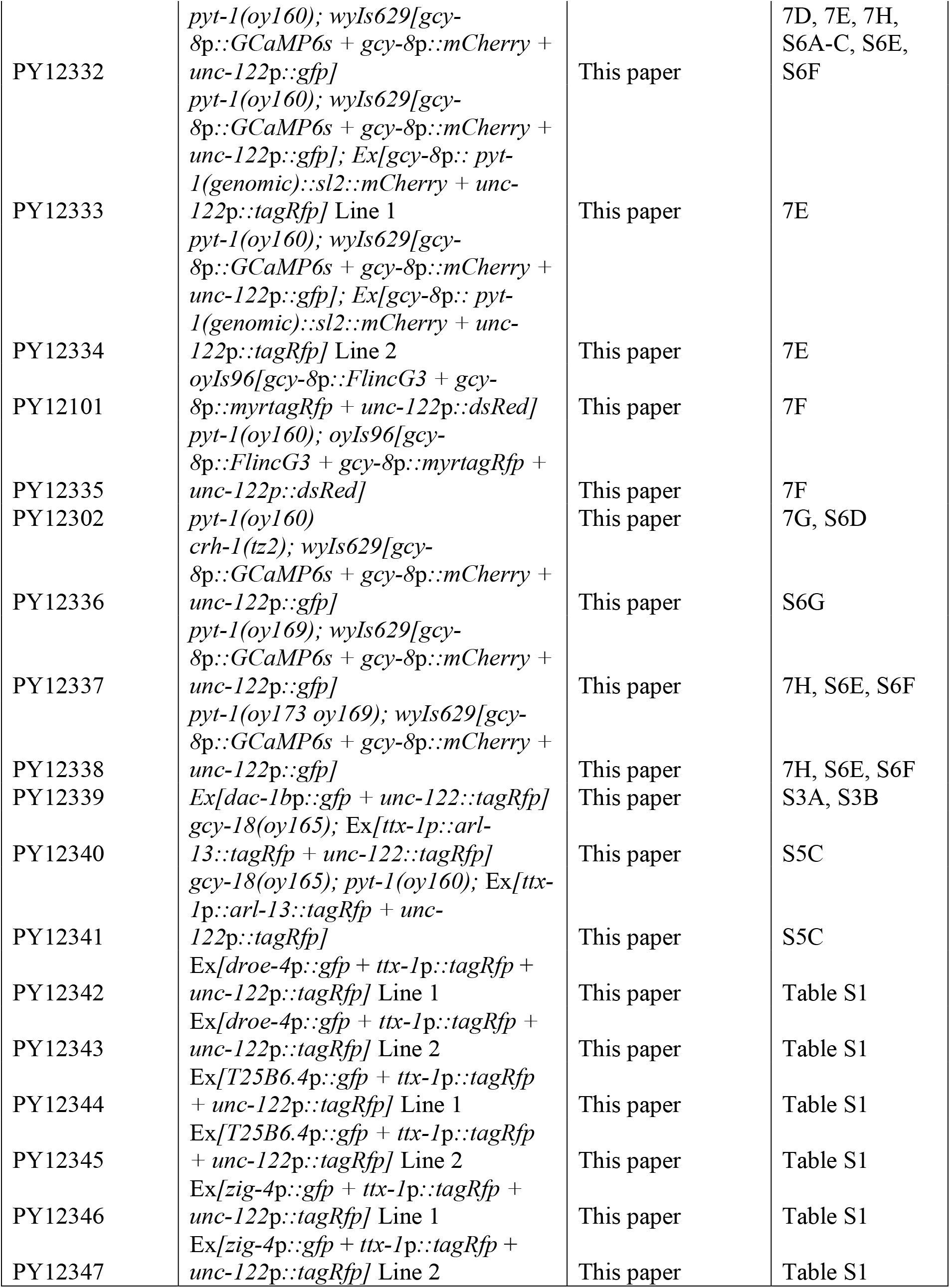
List of strains used in this work.

**Table S3.**
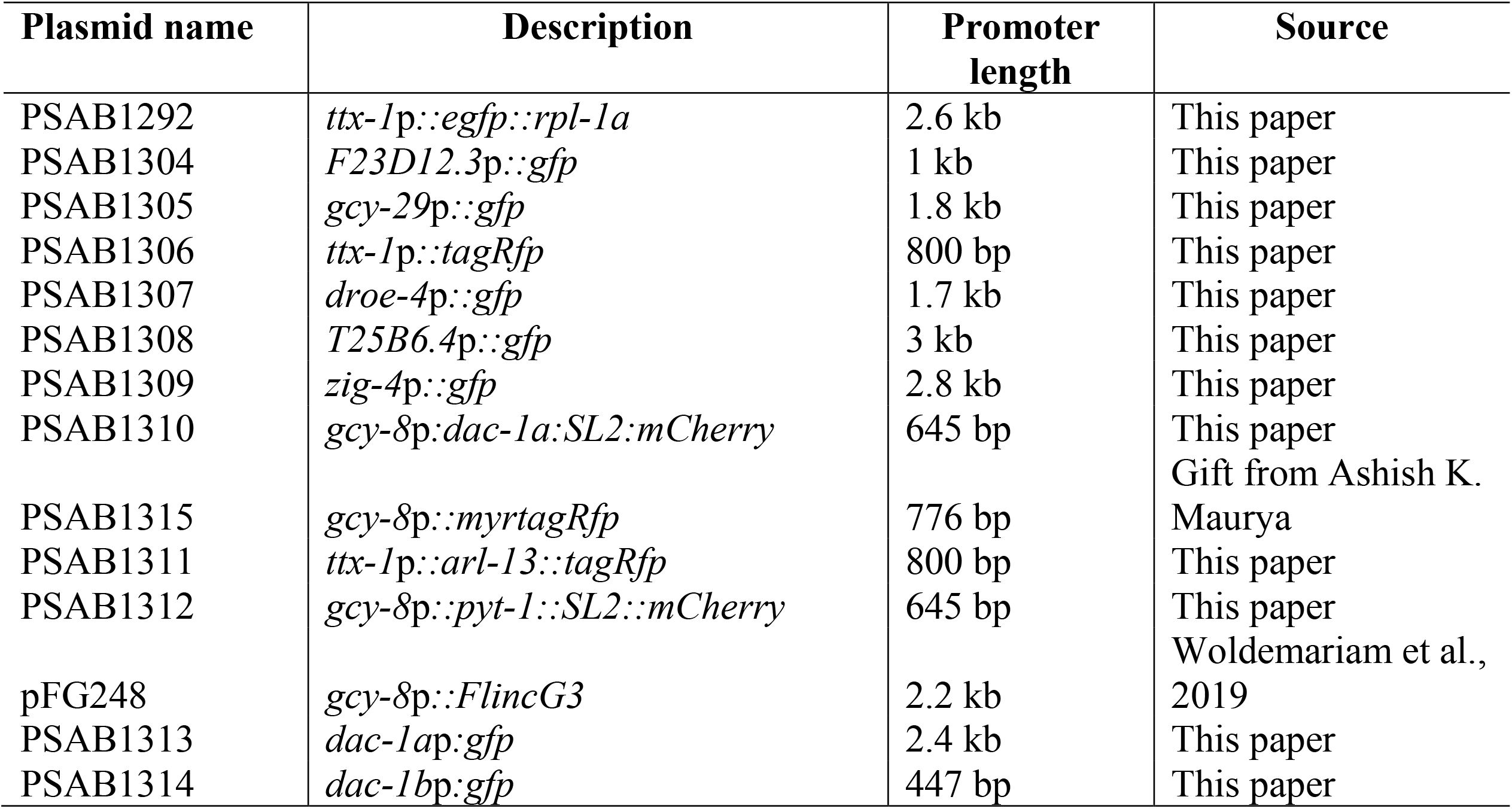
List of plasmids used in this work.

